# Consequences of directly- and indirectly-experienced heat stress in a mutagenic environment

**DOI:** 10.1101/2023.10.03.560724

**Authors:** Lauric Feugere, Claudio Silva De Freitas, Adam Bates, Kenneth B. Storey, Pedro Beltran-Alvarez, Katharina C. Wollenberg Valero

## Abstract

Climate change increases the frequency and duration of heat events. Negative effects of heat stress may be exacerbated through the action of social metabolites between aquatic animals. Whilst early life stages are vulnerable to stress-induced damage, they deploy cellular mechanisms to protect cells against mutagens such as ultraviolet rays (UV). Little is known about the fate of fish embryos which have experienced heat stress in a mutagenic environment. The present study exposed zebrafish embryos to one of three stress history treatments consisting of direct heat stress (TS+UV), the social context of heat stress via social metabolites (SM+UV), and their combination in TS+SM+UV before a UVB/UVA damage/repair assay. We measured phenotypic and transcriptomic responses to these treatments, and estimated mutational damage through DNA mutation frequencies and RNA integrity values. Compared to UV-treated controls (C+UV), the social context of heat stress history preceding the UV assay altered keratin and cell structuring-related pathways, associated with longer embryos with over-developed pericardia displaying behavioural hypoactivity. Relative to C+UV, direct heat stress history preceding UV exposure had a hormetic effect by stimulating the cellular stress response and facilitating DNA repair, which rescued embryos from subsequent UV damage and improved their apparent fitness. However, heat stress combined with social metabolites overwhelmed embryos in the UV assay, which annihilated the hormetic effect, introduced mutations, and lowered their apparent fitness. Whilst generated in the laboratory, these findings provide an important baseline for understanding the consequences of heat stress history in natural environments, where heat stress occurs within a social context.

**Highlights:**

- Heat stress had a hormetic effect against UV damage, by stimulating the heat shock response, antioxidants, and DNA repair.
- The heat hormetic effect protected and/or rescued embryos from UV damage by reducing single nucleotide variants observed in RNA, lowering malformations, and accelerating development.
- Heat-stressed embryos released social metabolites that initiated keratin, immune, and cellular structuring responses in receivers, in turn increasing body sizes but without reducing UV-induced malformations.
- Heat combined with social metabolites overwhelmed embryos in response to UV, reducing fitness-relevant performance.
- Heat stress during early embryogenesis led to differential fitness-relevant outcomes showing a nonlinear relationship with stress intensity.

**Summary statement:** Sublethal heat stress protects zebrafish embryos in a mutagenic environment, but this protective effect is lost when zebrafish embryos additionally stress each other via chemical cues.

**Graphical abstract:** 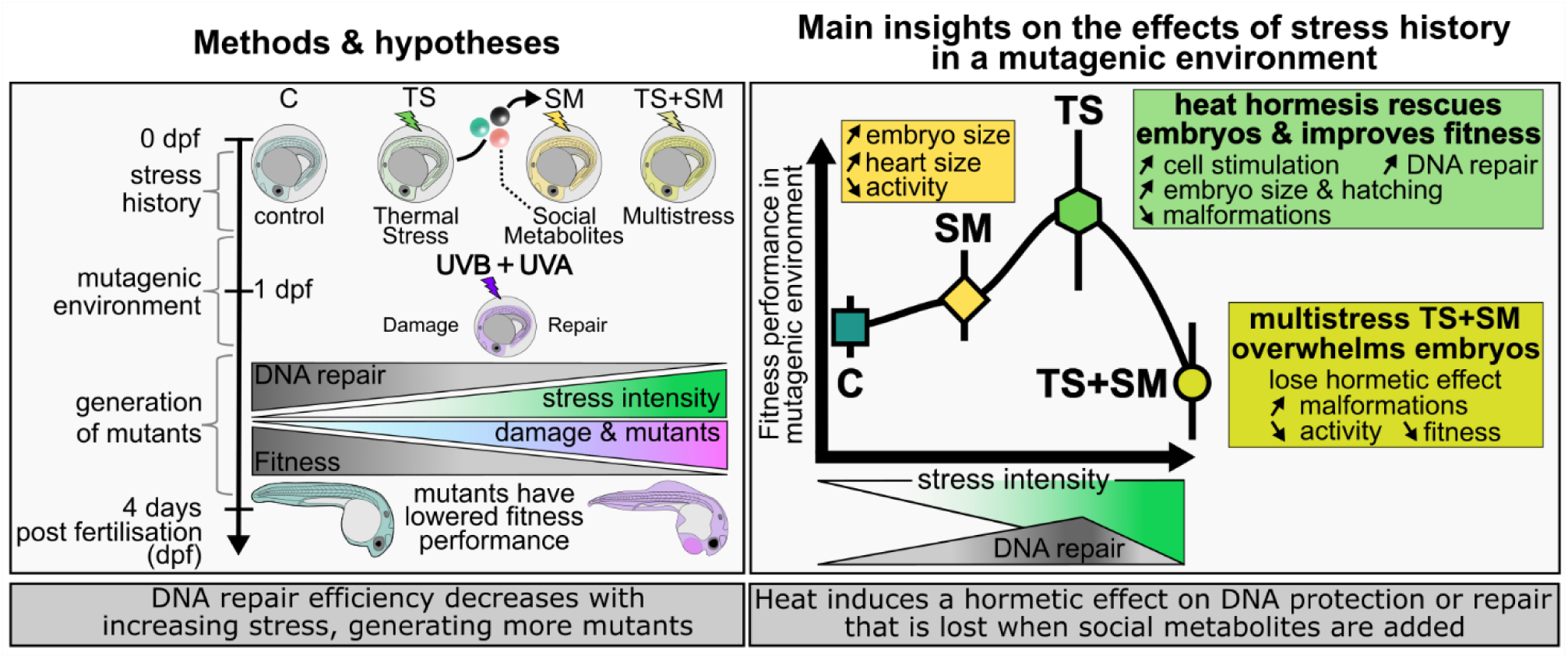

## Introduction

In an era of fast-paced climate change, thermal anomalies such as extreme temperatures are more frequent (IPCC, 2019; 2022). Such heat events may cause sharp declines in fitness in ectotherms (Jørgensen et al., 2022). Aquatic ectotherms, such as zebrafish (Engeszer et al., 2007), may spawn in shallow waters where offspring experience high temperatures and environmental mutagens such as ultraviolet radiation (UVR) from sunlight (Dahms and Lee, 2010). Moreover, heat-stressed aquatic animals may release “social metabolites” that alter the phenotype of conspecifics not directly experiencing heat stress (Feugere et al., 2021a; 2021b). Embryos developing in clutches may have amplified stress responses to heat through chemical communication, the consequences of which have received little focus. Early life stages developing outside the maternal body are most vulnerable to both UVR (Dahms and Lee, 2010; Zagarese and Williamson, 2001) and heat stress (Dahlke et al., 2020a; 2020b). Since early-life exposure to UVR lowers fitness in later stages (Ceccato et al., 2016; Lundsgaard et al., 2022), it is important to understand the combined effects of heat stress and environmental UVR on developing stages, as past studies found that abiotic stressors, such as high temperature, combined with UVR can lead to additive, synergistic, or antagonistic effects on the biology of fish (Alves and Agustí, 2020; Cramp et al., 2014; Icoglu Aksakal and Ciltas, 2018). For instance, heat may denature DNA repair proteins (Alton and Franklin, 2012; Lupu et al., 2004) and lower energy investment allocated to DNA repair (Berger et al., 2017). A lowered DNA repair capacity may increase DNA damage, which would affect fitness and survival through increased mutation rates. Prior studies mostly investigated heat simultaneous to UVR. However, stressors can have “latent carryover” effects persisting once the initial stressor is removed, which is an important yet overlooked avenue of research (Lundsgaard et al., 2023; O’Connor et al., 2010; O’Connor et al., 2014; Todgham and Stillman, 2013). The effects of UVR combined with temperature may be complex and depend on the timing of heat exposure (Hird et al., 2023). In this study, we investigate carryover effects, which we refer to as “stress history”, when encountering a subsequent mutagenic environment.

In natural environments, both ultraviolet A (UVA, 315-400 nm) and ultraviolet B (UVB, 280-315 nm) filter through the water column and cause molecular damage to lipids, proteins, and DNA (Alves and Agustí, 2020; Dahms and Lee, 2010; Rastogi et al., 2010). Most DNA damage is attributable to UVB, whilst UVA (along with blue light) activates photorepair mechanisms that directly reverse DNA lesions such as cyclobutane pyrimidine dimers (CPDs) and 6-4 photoproducts (6-4PPs; Banaś et al., 2020; Dong et al., 2008; Rastogi et al., 2010). Other repair mechanisms include nucleotide excision repair to substitute damaged nucleotides (Dahms and Lee, 2010; Griffiths et al., 1998; Rastogi et al., 2010), mismatch repair to remove mispaired bases (Jiricny, 2013) and homologous recombination and non-homologous end joining to repair double-strand breaks (Minten and Yu, 2019; Rastogi et al., 2010). Failure to mount such repair responses to UVR can impair immunity (Cramp et al., 2014; Salo et al., 2000), and scale up to whole-body damage such as altered behaviour, locomotion, increased malformation, and in turn, mortality (Alves and Agustí, 2020; Downie et al., 2023; Hurem et al., 2018). UVB may also induce inheritable epigenetic changes such as histone modifications, resulting in inherited resistant phenotypes (Reardon et al., 2022; Shen et al., 2017). The well-characterised cellular response to UVR can be harnessed to better understand the damage and repair systems following direct and indirect (via social metabolites) heat stress history. By limiting DNA repair capacity (Washington et al., 2003) and increasing reactive oxygen species (Xu et al., 2021), stress history may additionally increase the odds of spontaneous single-nucleotide variant (SNV) DNA mutations in subsequent UVR exposure (Sugiyama and Chen, 2019), which then can be observed in mRNA transcripts if they occur in transcribed, protein-coding genes (Castle et al., 2014).

Zebrafish (*Danio rerio*), even at embryonic stages, possess a competent repair system, involving genes in all repair pathways, to remove UVB-induced DNA lesions (Dong et al., 2007; Dong et al., 2008; Pei and Strauss, 2013; Sussman, 2007). Among these, blue light- and UVA-activated photolyases efficiently photorepair CPDs and 6-4PPs formed during UVB exposure, rescuing zebrafish embryos from UVB-induced morphological defects (Dong et al., 2007; Dong et al., 2008). We recently evidenced that heat, experienced both directly (sublethal heat stress) and indirectly (heat-induced social metabolites) stressed zebrafish embryos (Feugere et al., 2021b; 2023). We now ask whether this heat stress history affects DNA repair capacity in a mutagenic environment in terms of nucleic acid integrity and differential expression of repair-relevant genes, and, as a result, affects fitness-relevant outcomes (reaction to the novel stimuli light and touch) at the phenotypic level. We proposed that heat stress history, both directly and within a social context, could weaken the reparative response to UVR, and that this would be further disrupted by their combination, which would be observable through altered repair-related gene expression, decreased RNA integrity, and an increase in low-frequency nucleic acid mutations.

## Materials and Methods

### Zebrafish handling methods

Adult zebrafish (*Danio rerio*, AB strain) stocks were obtained from the University of Sheffield and the University of Cambridge, and were maintained at the University of Hull in a temperature-controlled room at 27°C with a 14:10 light:dark cycle. Fish were fed twice a day an alternate diet of mini-bloodworm, daphnia, and dried flakes. Breeding consisted of placing plastic trays filled with marbles and plastic plants, directly in the adult fish stock tanks, in the afternoon prior, and collecting zebrafish embryos at 10 am. Zebrafish embryos were collected using plastic pipettes and placed in cups with system water before being transferred to the laboratory. Zebrafish embryos were cleaned in fresh E3 embryo medium (Cold Spring Harbor Laboratory Press, 2011) to remove organic matter. Next, zebrafish embryos were immersed in a small tea strainer with gentle swirling in small petri dishes to carry out a bleaching protocol of 3 min in each of the following baths: (1) bleaching medium, (2) embryo medium, (3) bleaching medium, (4) embryo medium, and (5) embryo medium. Bleaching medium contained 0.004% bleach in E3 embryo medium after diluting 10-13% active sodium chloride in embryo medium. Only viable embryos (at least 2-cells stage and no more than at the high stage i.e. 3.3 hours post fertilisation, with no visible deformation) were carefully pipetted using a glass Pasteur pipette into 0.2 mL PCR wells pre-filled with the experimental medium. All experiments were approved by the Ethics committee of the University of Hull (FEC_2019_194 Amendment 1). Zebrafish stages are expressed in hours post fertilisation (hpf) and days post fertilisation (dpf) according to Kimmel et al. (1995).

### Experimental design

One hour after collection and immediately after bleaching protocols, zebrafish embryos were individually exposed for 24 hours to experimental stress history treatments, starting at 11 am during the cleavage period (2 to 3.3 hpf) until 11 am on the next day when embryos were 1 dpf. Zebrafish embryos were exposed to a two-way factorial design of two temperature protocols (control *vs.* sublethal heat stress) and two types of embryo medium (fresh medium *vs.* medium containing social metabolites). Embryos were housed individually in a thermal cycler with closed lid, in 0.2 mL PCR tubes containing 200 µL of their experimental medium and temperature. Here, we define “stressors” as environmental factors that alter homeostasis, which can be observed by measuring the behavioural and physiological adaptive responses aiming at restoring homeostasis (Barton, 2002; Barton and Iwama, 1991; Chrousos, 1998; Chrousos, 2009). We have recently shown that repeated exposure to sublethal temperature of 32°C is stressful to zebrafish embryos, posing the framework to study the effects of both direct heat stress history and the social context of heat stress propagated via chemical cues (Feugere et al., 2023). The fresh medium was free of any metabolites whilst the stress medium contained “social metabolites” released by heat-stressed conspecific donors as described in Feugere et al. (2023). We have previously shown that three types of media containing social metabolites from stressed animals, control metabolites from unstressed animals, or no metabolites induce distinct phenotypes and react differently when combined with abiotic stressors (Feugere et al., 2023). For this reason, we opted for a control free of metabolites to be able to use a factorial design of single and combined treatments with heat stress and social metabolites. The factorial design yielded the following condition: control (C) or one of three “stress history” treatments, namely, thermal stress (TS), social metabolites (SM), or a combined treatment (TS+SM). The treatments SM, TS, and TS+SM were considered heat stress history treatments where TS is a direct heat stress and SM is the social context of heat stress. However, the trade-off from this design was that in this study we cannot discern whether the effect of social metabolites stem from regular social cues or specific ones induced by heat stress. After 24 hours of exposure, 1-dpf embryos were removed from their treatments and exposed to a UVB/UVA damage/repair assay (see Fig. 1; *Experimental UVR exposure,* below).

**Figure 1.**
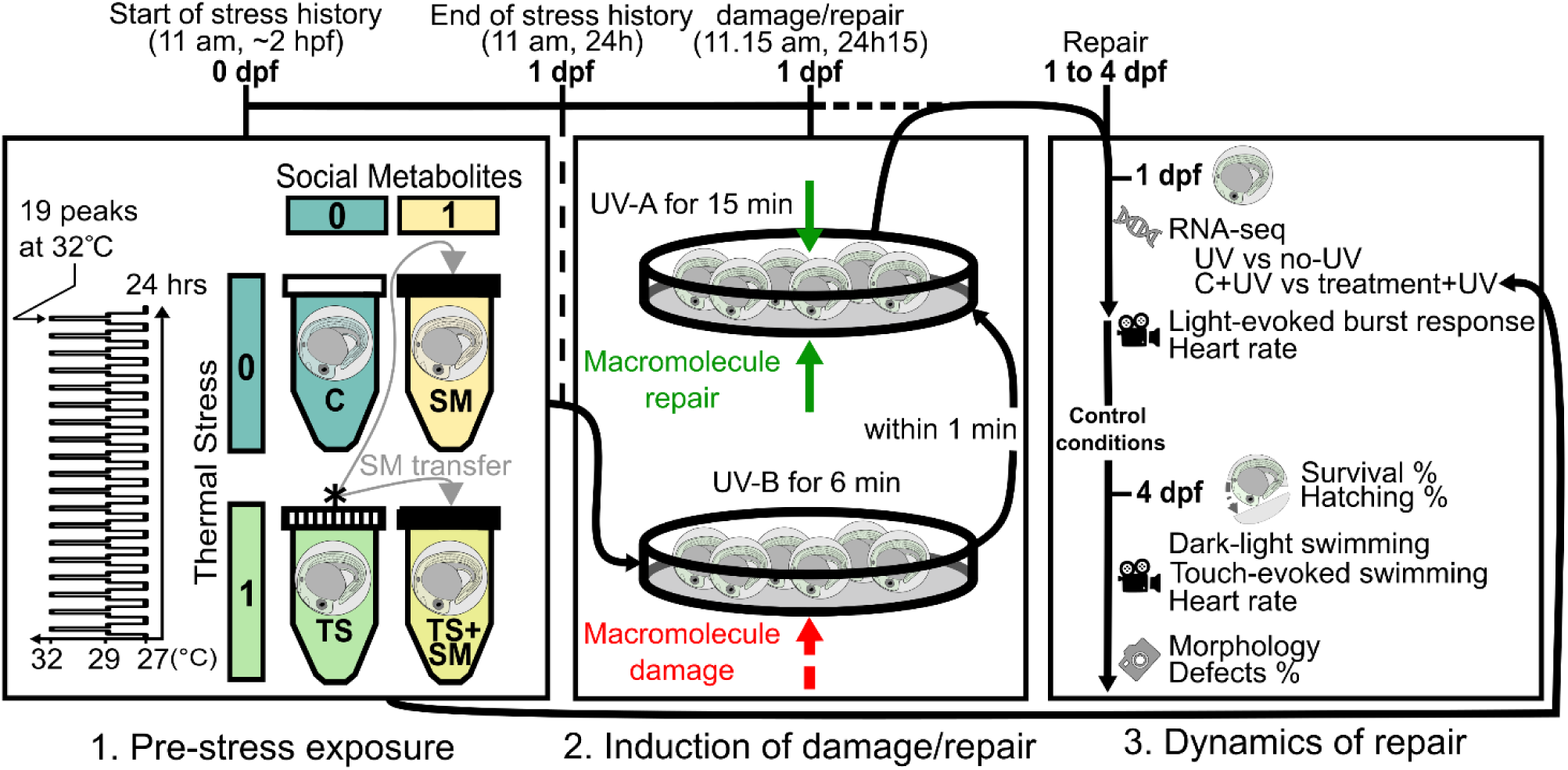
Scheme of experimental design. Left: zebrafish embryos were exposed individually to heat stress history conditions of repeated thermal stress (see diagram of heat peaks) or social metabolites induced by it from 2 hrs post fertilisation (hpf) to 1 day post fertilisation (dpf). Stress history treatments were: control (C), thermal stress (TS), social metabolites (SM), social metabolites in thermal stress (TS+SM). Grey arrows show metabolite transfer from donors (*, hatched tube caps) to receiver embryos (black tube caps). Middle: At 1 day post fertilisation (dpf), embryos experienced successive macromolecule damage through ultraviolet B (UVB, short-dash red arrow) and repair through ultraviolet A (UVA, plain green arrows). Right: zebrafish embryos were either immediately sampled for RNA-sequencing at 1 dpf (with or without UV) or incubated in control conditions for phenotypic analyses until 4 dpf. Endpoints were RNA-sequencing (DNA symbol), morphology (camera), hatching (opened chorion), behaviour (video recorder), and survival.

This assay induces deleterious mutations through UVB exposure followed by UVA-induced DNA repair. UVR treatments were labelled C+UV, TS+UV, SM+UV, and TS+SM+UV. Zebrafish embryos were sampled both before (data from Feugere et al., 2023) and immediately after UVR exposure at 1 dpf for RNA-sequencing. Another set of embryos was maintained under control conditions (i.e. 27°C, in a thermal cycler with closed lid, in 0.2 mL tubes containing 200 µL of fresh medium) after UVR exposure until 4 dpf for phenotypic analyses. Embryos were humanely euthanized before 5 dpf through snap-freezing at −80℃.

### Temperature protocols

Temperature protocols were either (i) a control constant temperature of 27°C, or (ii) a thermal stress consisting of 19 cycles of temperature fluctuations between 27, 29, 32, 29, and 27°C, with each step maintained for 15 min. Whilst deviating from realistic heatwaves, this thermal stress protocol aimed to expose embryos to a maximum repeat of sublethal temperature of 32°C and mimic +5°C heat events at crucial times of embryonic development. Whilst less-plastic laboratory-adapted laboratory zebrafish strains may have different response compared to their wild counterparts (Morgan et al. 2022; Vossen et al. 2020), the general thermal tolerance of domesticated zebrafish may not change drastically and they can be used to address certain ecological questions (Morgan et al. 2019). Temperature protocols were maintained using thermal cycles with a closed lid, causing constant darkness that delays the normal zebrafish embryonic development (Villamizar et al., 2014).

### Embryo medium

A 60× stock solution of embryo medium contained 34.8 g NaCl, 1.6 g KCl, 5.8 g CaCl_2_·2H_2_O, and 9.78 g MgCl_2_·6H_2_O in 2 litres of ultrapure water (Cold Spring Harbor Laboratory Press, 2011). After adjusting to pH = 7.2 with NaOH, the 60× stock solution was autoclaved and stored at 4°C. The working solution of 1× E3 embryo medium, stored at 27°C and replaced at least weekly, was prepared by 1:60 dilution of the 60× stock in ultrapure water. Social metabolites were obtained by pooling 180 µL of medium per heat-stressed donor’s well into a Falcon tube and immediately transferring 200 µL of the pooled stress medium into social metabolite receivers’ wells. Fresh social metabolites for SM and TS+SM were obtained from donors exposed to TS on the day prior to limit their molecular degradation.

### Experimental UVR exposure

At the end of the 24-hour-long exposure to C, SM, TS, and TS+SM, survival rates were monitored and viable embryos were placed into small petri dishes (35 mm in diameter) in their incubation media for each treatment, and were subjected to UVR in the UVB/UVA damage/repair assay. The methods were adapted from Dong et al. (2007) and optimised for allowing >75% survival at 4 dpf to measure differences in phenotypes between treatments (see “*Optimisation of UVR exposure”* in Supplementary Information). We used this sequential exposure to UVB and UVA, instead of simultaneous exposure, to ensure that UVB induced macromolecule damage that could be repaired through UVA, allowing us to study the changes in DNA repair capacity depending on stress history. The UVB/UVA damage/repair assay was first optimised on control embryos so that UVA rescued embryos from UVB-induced defects with both no excess mortality and a significant increase in healthy embryos. For the experimental exposure to UVB/UVA, embryos were placed in small petri dishes (35 mm in diameter) with ∼5 mL of fresh embryo medium. Zebrafish embryos within their chorion were centred in their Petri dish and exposed at 27°C in darkness for 6 min on top of a UV transilluminator (Syngene Ltd., GelVue GVM20) equipped with 6 UVB bulbs (peak of 306 nm, UVB, 8 watts lamp wattage, 1.6 watt ultraviolet output, Sankyo Denki G8T5E). Next, zebrafish embryos were immediately transferred (within 1 min) to the UVA setup which consisted of a 15-min exposure at 27°C in a ‘sandwich’ system of PCR handle stations (365 nm UVA, 4W UVA output, Spectroline ENF-240C/FE) comprised of one bottom-up (direct contact against the bottom of the Petri dish) and one on top-down (11 mm away from embryos due to the height of the petri dish) UVA sources. The UVA setup was placed in the laboratory under both visible light from bulbs and natural sunlight which also excite DNA repair enzymes such as photolyases (Steindal and Whitmore, 2020). The UV irradiance could not be measured in the laboratory. The chosen timing (6 min + 15 min) aimed to allow the expression of DNA repair mechanisms at the time of embryo sampling for RNA-sequencing (within 15 min after the end of the UVB/UVA exposure).

### RNA-sequencing and analysis

RNA sequencing was performed using the same methods and sequenced at the same time as the samples described in Feugere et al. (2023). At the end of the stress history treatments and immediately after UVR assays, embryos still in their chorion were pooled by groups of 20 individually-exposed embryos and immediately humanely euthanised by snap-freezing at −80°C. Sample sizes for RNA-sequencing were n = 3 biological replicate pools for each treatment, each representing the average gene expression of 20 embryos per replicate and 60 embryos per treatment. Pooling was determined as the best method to increase statistical power whilst reducing noise from individual variation in gene expression (Takele Assefa et al., 2020). The experiment was replicated on different clutches of embryos until enough samples were generated for sequencing. Samples were randomised, at the time of RNA extraction, for blind sample preparation and RNA-sequencing. Total RNA was extracted via the TRIzol method followed by a DNAse I digestion step (#10792877, Invitrogen™ TURBO DNA-free™ Kit) and a sodium acetate clean-up. Total RNA samples were assessed for quality, cDNA libraries were prepared with the TruSeq stranded mRNA kit, and sequenced by Illumina NovaSeq 6000, 50PE sequencing at the Edinburgh Genomics facility. FASTQC v0.11.9 (https://www.bioinformatics.babraham.ac.uk/projects/fastqc/) was used to assess read quality before and after filtering and trimming using *fastp* v0.23.1 (Chen et al., 2018). Reads from UV-exposed conditions were deduplicated using *fastp* after PCR-biased duplication was detected using the R package *DupRadar* v1.18.0 (Sayols et al., 2016). The splice-aware mapper *STAR* v2.6.1 (Dobin et al., 2013) served to align and count reads to the reference zebrafish genome (primary genome assembly GRCz11.104). In addition to the UV-samples generated in this contribution (available under accession number GSE223685 on NCBI’s Gene Expression Omnibus database), corresponding RNA sequence data from non-UV treatments (C, TS, SM, TS+SM, available under accession number GSE220546 on NCBI’s Gene Expression Omnibus database) were taken from Feugere et al. (2023).

*DESeq2* v1.28.1 (Love et al., 2014) was used for differential expression analyses in R v4.0.2 (R Core Team, 2020) using *Bioconductor* v3.11 (Morgan, 2021). Taking advantage of a three-way factorial design of heat × social metabolites × UVR, we tested for effects of stress history on the response to UVR. For this, all three treatments SM+UV, TS+UV, and TS+SM+UV were compared to C+UV (Dataset S1). The effect of the combined stressors (direct heat and social metabolites) was explored by comparing SM+UV and TS+UV to TS+SM+UV (Dataset S1). If heat and social metabolites do not have an effect in interaction, one would expect that comparing TS+SM+UV to SM+UV would be equivalent to comparing TS+UV to C+UV for the heat stress term, whereas comparing TS+SM+UV to TS+UV would be the same as comparing SM+UV to C+UV for the social metabolites term. Conversely, relative to TS+SM *vs*. C, if different patterns are found when comparing the term TS+SM to the single treatment terms (SM or TS), this indicates that there is an interaction effect of TS and SM. The transcriptomic response to UVR was then investigated by comparing each UV versus no-UV treatment pair, i.e., C *vs.* C+UV, SM *vs.* SM+UV, TS *vs.* TS+UV, and TS+SM *vs.* TS+SM+UV (Dataset S1). Third, the magnitude of change (absolute log fold change, LFC) and gene expression ratios in response to UVR were compared in a two-way design of heat × social metabolites across C+UV, SM+UV, TS+UV, and TS+SM+UV. This analysis was repeated with Gene Ontology (GO)-term wide p-value adjustments within subsets of genes from four candidate pathways that may be involved in facilitating recovery from UVB: “*DNA repair”* (GO:0006281), “*protein folding”* (GO:0006457), “*macromolecule methylation”* (GO:0043414), and “*regulation of gene expression, epigenetic*” (GO:0040029). Principal Component Analyses of regularised logarithmic (*rlog*) transformed data were explored using the *prcomp* R function from the *stats* package (R Core Team, 2020). The differentially expressed genes (DEGs) were visualised as volcano plots with the *EnhancedVolcano* R package v1.6.0 (Blighe et al., 2020). DEGs were considered significant when p-adj < 0.05 and |FC| ≥ 1.5 (|LFC| > 0.58). Gene identifiers were cross-database-converted using *biomaRt* R package v2.44.4 (Durinck et al., 2005; Durinck et al., 2009) and mapped to GO terms using *topGO* v2.40 (Alexa and Rahnenfuhrer, 2020) and *org.Dr.eg.db* v3.11.4 (Carlson, 2020). *GOstat* v2.54.0 (Falcon and Gentleman, 2007) (hypergeometric test), *clusterProfiler* v3.16.1 (Yu et al., 2012), and *ReactomePA* 1.32.0 (Yu and He, 2016) R packages served to identify functionally enriched GO terms, Kyoto Encyclopedia of Genes and Genomes (KEGG) pathways, and Reactome pathways, respectively. Functional terms were considered significantly enriched when p-adj < 0.1 and gene count ≥ 2.

### Phenotypic experiments

Phenotypic measurements consisted of imaging 4-dpf larvae for morphometric measurements, and videoing for heart rate and light-induced startle response (at 1 dpf), and light- and touch-evoked swimming (at 4 dpf). Larvae presenting less defects were considered to have a higher apparent fitness. Startle, light- and touch-evoked responses are standard procedures to measure fear, anxiety, and indicate underlying impairments to motor and sensory functions (Basnet et al., 2019; Beppi et al., 2021; Colwill and Creton, 2011; Golla et al., 2020). Zebrafish larvae may increase swimming in response to visual or tactile stimuli (Colwill and Creton, 2011). We consider the stimuli-evoked responses to simulate a risk environment that can provide insights on the effect of stress history on the apparent fitness of the larvae. UV-exposed larvae had mutant phenotypes that lowered their responsiveness to risk simulated by touch and light stimuli. Therefore, embryos that moved more at 1 dpf and larvae that showed more pronounced escape responses (i.e. higher behavioural activities) at 4 dpf were considered to have a higher apparent fitness by avoiding the simulated risk.

After treatments and immediately after UVB+A exposure, 1-dpf zebrafish embryos were moved per treatment groups into small glass dishes for video recording of startle responses. Zebrafish embryos within their chorion were videoed for burst activity (for 15 s under the stereomicroscope after switching the underneath light source to 50% intensity to induce the startle response) by batches of 4-5 embryos in small glass wells with fresh embryo medium, with treatment alternation and blind to the experimenter. Zebrafish embryos were then placed in individual wells in control conditions (i.e. in 200 of µL of fresh embryo medium for incubation in darkness at 27°C). These embryos were uniquely identified and were re-used for more phenotypic measurements until 4 dpf. After 31 hours (6 hours after UVR treatment), each chorionated embryo was removed from its individual well, carefully placed using a plastic pipette into a glass watch in a small volume of fresh embryo medium and videoed for 15 s using the stereomicroscope for heart rate monitoring. Zebrafish embryos were then returned to their wells at which point 180 µL of their medium was replaced with 180 µL of fresh embryo medium. Embryos were also monitored for survival (by transparency and presence of a heart beat under the stereomicroscope) and hatching at 2 and 3 dpf at 11 am at which point 180 µL of their medium were replaced with 180 µL of fresh embryo medium. At 4 dpf, zebrafish embryos that did not naturally hatch were manually dechorionated using fine forceps (Dumont n°5) before being returned to their individual wells. Embryos were allowed to recover from dechorionation in control conditions for 2 hours. At 5 pm at 4 dpf (102 hours of experimental exposure), zebrafish embryos were videoed for swimming assays using a Canon 1200D camera set at 25 frames per second. Zebrafish embryos were placed individually in 6-well plates filled with 3 mL of fresh embryo medium preheated at 27°C. Zebrafish embryos were left for five minutes in darkness and immediately videoed for two minutes once artificial light was applied to induce a startle-like response (dark-light swimming behaviour). Next, the embryos were videoed for a touch-evoked swimming behaviour which consisted of three consecutive gentle touches on the head, every ten seconds, using an inoculating needle. The embryo medium was changed between each embryo. At the end of the swimming assay, after 105 hours of treatment (8 pm on day 4), zebrafish embryos without their chorion were carefully moved on a microscope slide labelled with a micrometre for imaging under the stereomicroscope. Embryos were carefully moved into a glass watch in a small volume of fresh embryo medium and videoed for 15 s using the stereomicroscope to acquire 4-dpf heart rate videos.

All metrics of phenotypic measurements (hatching, survival, morphometry and behaviour) were observed from embryos in an experiment performed once on a single batch of zebrafish embryos with the following sample sizes: n = 28 (C+UV), n = 22 (SM+UV), n = 20 (TS+UV), and n = 25 (TS+SM+UV). Sample size was determined by the number of available viable embryos from a clutch at the time of testing, evenly spread across conditions within the constraints of available social metabolite medium available from the day prior. In all phenotypic analyses, treatments were alternated to limit time effect on swimming behaviours, and blind to the experimenter to limit experimental bias. Images and videos were randomised for blind analysis using Bulk Utility Rename v3.4. Videos of startle response at 1 dpf and heart rates (heart beat per minute) at 1 and 4 dpf were analysed using Danioscope (Noldus). The data of burst activity percentage (percentage of the time, from the total measurement duration, that the embryo was moving) and burst count (number of times the embryo moved) per minute were retrieved from the startle response videos. Because automatic data acquisition softwares did not correctly track larvae for our videos of 4-dpf swimming, these were analysed manually (i.e. by tracking the position of the head of larvae at each frame) using KINOVEA v0.9.5 (Charmant and Contributors, 2021). We obtained binary moving data (i.e. whether the larvae swam in response to stimuli), distance swam (in cm) and time the larvae was swimming (in s) data from which speed (in cm/s) was calculated. The number of times a larva accelerated and decelerated until it stopped swimming was counted and dubbed “swimming burst count”. Swimming data were obtained for the entire 2-min dark-light assay videos and following each of the three stimuli in the touch-evoked swimming assay. The total distance swam, average speed, and total time moving over the three touches were used in the statistics. Images were analysed using ImageJ v1.53e (Schneider et al., 2012) after calibration to a micrometre scale to measure lengths (mm) of several morphological characters: whole-body, eye, yolk ball, yolk extension lengths; and pericardial and tail widths.

### Minor Allele Frequency comparison

To assess nucleic acid damage induced by stress history treatments and UVR at the sequence level, Minor Allele Frequencies were called from transcriptomes with the somatic small-variant caller *Strelka2* against the zebrafish reference genome assembly (GRCz11.104). While variant calling from RNA-seq is not a true representation of the amount of genomic variants, it nonetheless allows for comparisons of expressed alleles between treatments (Castle et al., 2014). Transcriptomes were compared (i) for each UV-*vs.* no-UV treatment (C+UV *vs.* C, TS+UV vs TS, SM + UV *vs.* SM, TS+SM+UV *vs.* TS+SM) and (ii) between UV-treated control and stress history treatments (C+UV *vs.* TS+UV, C+UV *vs.* SM+UV, C+UV *vs.* TS+SM+UV) We discarded any resulting In/Del calls as these could have resulted from faulty transcript overlap during alignment, and filtered any resulting single nucleotide variants (SNV) for high EVS (Empirical variant scores). As per developer’s suggestions, variants with the Low Depth filter flag were included, as UVR can induce mutations in as little as one RNA transcript in each sample’s transcriptome. Since a pool of 20 diploid embryos per RNA-seq sample has 40 alleles at any given locus, one SNV in one of these embryos’ DNA would lead to a MAF of 0.025, while further sporadic mutations at RNA level would likely fall below this threshold. Therefore, we expected that any differences in MAF distribution between treatments in the 0.025-0.1 range and ideally at a frequency of 0.025, would be indicative of UV- or treatment-induced spontaneous SNV mutations in DNA. Differences in allele frequency distributions were assessed with Kolmogorov-Smirnov two-sample tests.

### Statistical analyses

All statistical analyses were performed in R v4.0.2 (R Core Team, 2020) and graphs mainly drawn with the *ggplot2* R package (Wickham, 2016). Gene expression binary data were analysed with chi-squared tests from *rstatix* v0.7.0 (Kassambara, 2021) using *chisq_test* for model terms and *pairwise_chisq_gof_test* for pairwise *post-hoc* treatment comparisons. Gene expression linear models were represented with *ggscatter* from *ggpubr* v0.4.0 (Kassambara, 2020). Binary phenotypic data were analysed using generalised linear models with binomial data. Parametric numerical data were analysed using two-way ANOVAs followed by pairwise post-hoc tests using *emmeans* v1.7.3 (Lenth, 2022), after transformation from *BestNormalize* v1.8.2 (Peterson, 2021; Peterson and Cavanaugh, 2020) where necessary and possible. Model assumptions were verified using Breusch-Pagan tests for heteroskedasticity from *lmtest* v0.9-38 (Zeileis and Hothorn) and Shapiro-Wilks tests for normality of residuals. Non-parametric numerical data were analysed using Scheirer-Ray-Hare tests followed by pairwise Wilcoxon-Mann-Whitney *post-hoc* tests from *rcompanion* v2.4.1 (Mangiafico, 2021). Since changes in morphometry (e.g. embryo size) were outcomes of treatments, we did not include them as covariates but instead tested for correlation statistics between phenotypic variables using *corrplot* v0.92 (Wei and Simko, 2021). PERMANOVAs were performed with the *adonis* function from *vegan* v2.5-7 (Oksanen et al., 2020) followed by pairwise PERMANOVAs using the *pairwise.adonis* function from *pairwiseAdonis* v0.4 (Martinez Arbizu, 2017). All tests are two-tailed and multiple comparisons were adjusted for False Discovery Rates.

As the treatments were optimised to be sublethal to the embryos and the observation period remained within the four days of development, we could not use mortality and reproductive success as proxies for fitness. Solely based on one behaviour, one cannot make assumptions about the consequences for the “apparent fitness” of the animal (Lind and Cresswell, 2005). However, combined with the different observed response variables (morphology, defects, several behaviours), we can build a better inference about the fitness of the embryos. Morphology can indeed be linked to fitness (Arnold, 1983; Lande and Arnold, 1983) although we acknowledge that there may be limits to linking morphology and behaviour to fitness (Cocciardi et al., 2021; Higginson, 2020) (preprints). Therefore, we aimed to represent the overall whole-body phenotypic response, based on all morphometric and behavioural data, as a proxy for the apparent fitness of the larvae. The statistical analyses of individual variables suggested that there was a nonlinear relationship between the fitness-relevant performance in response to UVR and the intensity of stress history is nonlinear. To verify this hypothesis, another set of analyses was conducted by performing separate multivariate analyses (PERMANOVAs) of the data at 1 dpf and 4 dpf to account for age-specific responses to UVR. For this purpose, Principal Component Analyses (PCA) of 1-dpf and 4-dpf data were conducted using *prcomp*. Multivariate data at 1 dpf included two variables: burst activity percentage and burst count per minute. Multivariate data at 4 dpf included 14 variables: number of burst events, percentage of time moving, speed, total distance, percentage of moving larvae in the dark-light assay; total distance moved, total time moved, and speed in the touch-evoked assay; as well pericardial edema, embryo length, embryo-size normalised pericardial width, embryo size-normalised eye length, hatching, and the inverted defect score. Multivariate data analyses were conducted for all complete cases, with missing cases being dependent (not pre-established) on survival. Fitness-relevant scores were the PCA scores from the first principal components (PC1) of the multivariate data, which corresponded to 70.7% and 40.3% of the variance contribution at 1 and 4 dpf, respectively. Fitness-relevant scores were re-coded prior to running multivariate analyses so that higher values indicated a higher apparent fitness (i.e. moving more and having fewer mutant phenotypes, as explained above). Fitness-relevant scores were compared between treatments using an ANOVA to summarise patterns within the whole of the data. Multivariate data were visualised using *ggbiplot* v0.55 (Vu, 2011). Model terms were visualised using the *plot_model* function from *sjPlot* v2.8.10 (Lüdecke, 2021) and trends were represented by smoothed *loess* lines using *stat_smooth* from *ggplot2*. We reported our results with the language of evidence when the evidence is weak (i.e., a “trend”, 0.1 < p < 0.05; Muff et al., 2022). Effect sizes were interpreted according to Sawilowsky (2009) as tiny (|d| < 0.1), very small (|d| > 0.1), small (|d| > 0.2), medium (|d| > 0.5), large (|d| > 0.8), very large (|d| > 1.20), and huge (|d| > 2.0).

## Results

### A common response to UVR existed in all treatments

We hypothesised that heat stress history amplifies the negative effects of UVR, which we tested through characterising the transcriptomic response to UVR (all statistical analyses of transcriptomic data are detailed in Dataset S1). The response to UVR was analysed by comparing each treatment followed by UVR against its non-UV counterpart (C *vs.* C+UV, SM *vs.* SM+UV, TS *vs.* TS+UV, and TS+SM *vs.* TS+SM+UV, Figs 2, S1).

**Figure 2.**
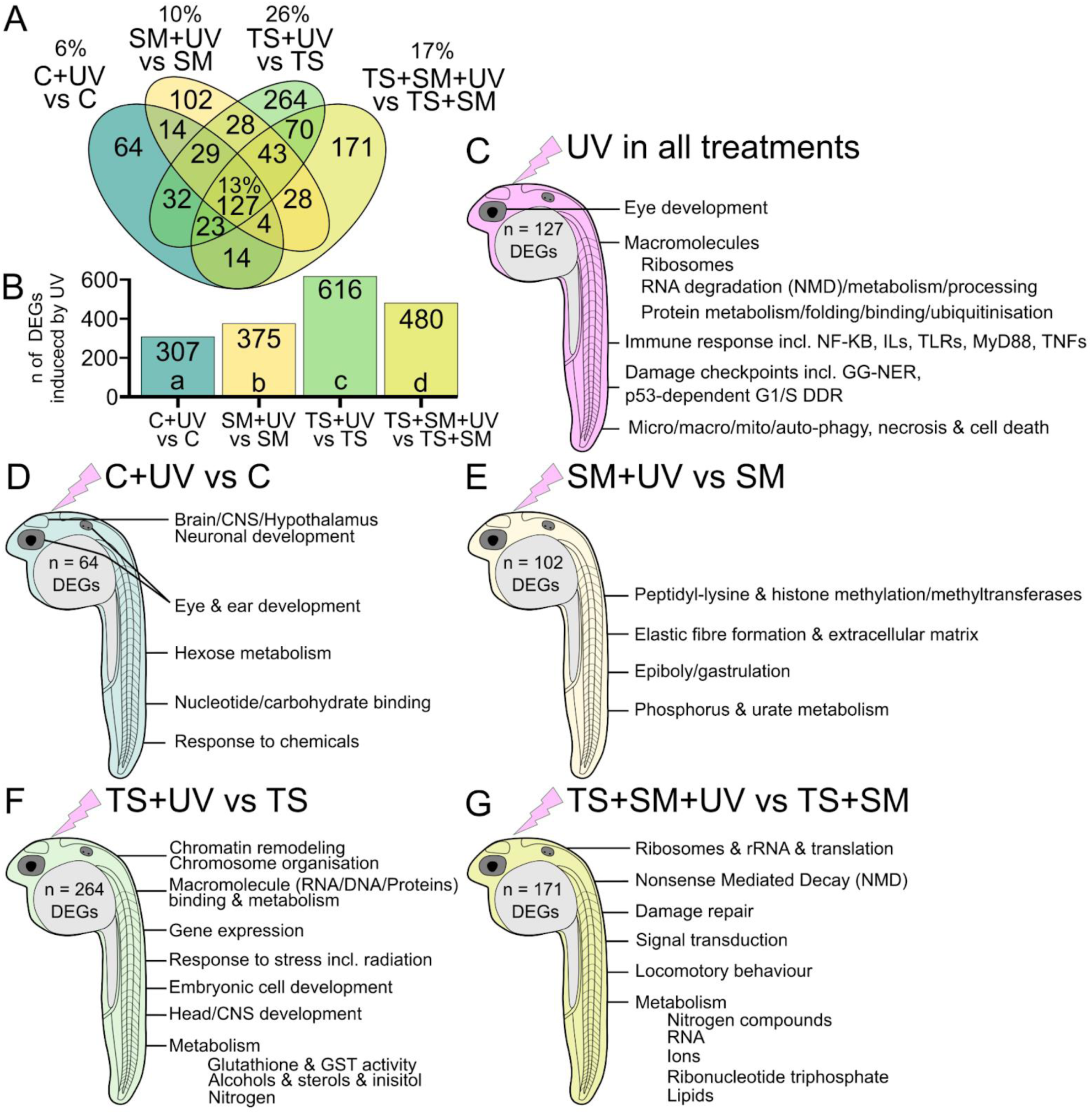
Heat stress history potentiates the transcriptomic response to UV exposure and induces a unique transcriptomic response to UVR with different biological signatures. Transcriptomic signature of each of the four UV treatments (C, SM, TS, TS+SM) compared to their non-UV pairs. A) Venn diagram of significantly altered genes in response to UVR. Functional enrichment of the genes (B) shared by all four treatments, or (C) unique to C+UV *vs.* C, (D) unique to SM+UV *vs.* SM, (E) unique to TS+UV *vs.* TS, and (F) unique to TS+SM+UV *vs.* TS+SM. Enrichments show the top enriched terms (with highest gene count/term) of Biological Processes, Molecular Functions, KEGG and Reactome pathways sorted by decreasing significance from lists of significant genes (p-adj < 0.05 and |FC| > 1.5). Treatments were C: control in fresh medium at 27°C, SM: social metabolites at 27°C, TS: fresh medium in thermal stress, TS+SM: social metabolites in thermal stress, all compared to their non-UV pairs. Sample size is n = 3 biological replicate pools of 20 embryos per treatment.

There were 127 differentially expressed genes (DEGs) in all treatments in response to UVR, with similar directionality and expression levels (Figs 2A, S1A). Shared genes were ascribed to macromolecule metabolic processes, nonsense-mediated decay (NMD), damage checkpoints, immune response, apoptosis/autophagy, and organ development (Figs 2C, S1B). UVR halved the transcription of 30 ribosomal protein subunits under all four treatments and altered the expression of heat shock proteins (*hsp70.2*, *hsp70l, hsp90ab1,* and *hspa8*) involved in the heat shock response (HSR).

### Different treatments initiated unique signatures and stress history potentiated the response to UVR

We confirmed that in a mutagenic environment, both social metabolites and heat functioned as stressors that altered major transcriptomic pathways after UVR exposure (Figs 3A,B,D,E, S1C-G).

**Figure 3.**
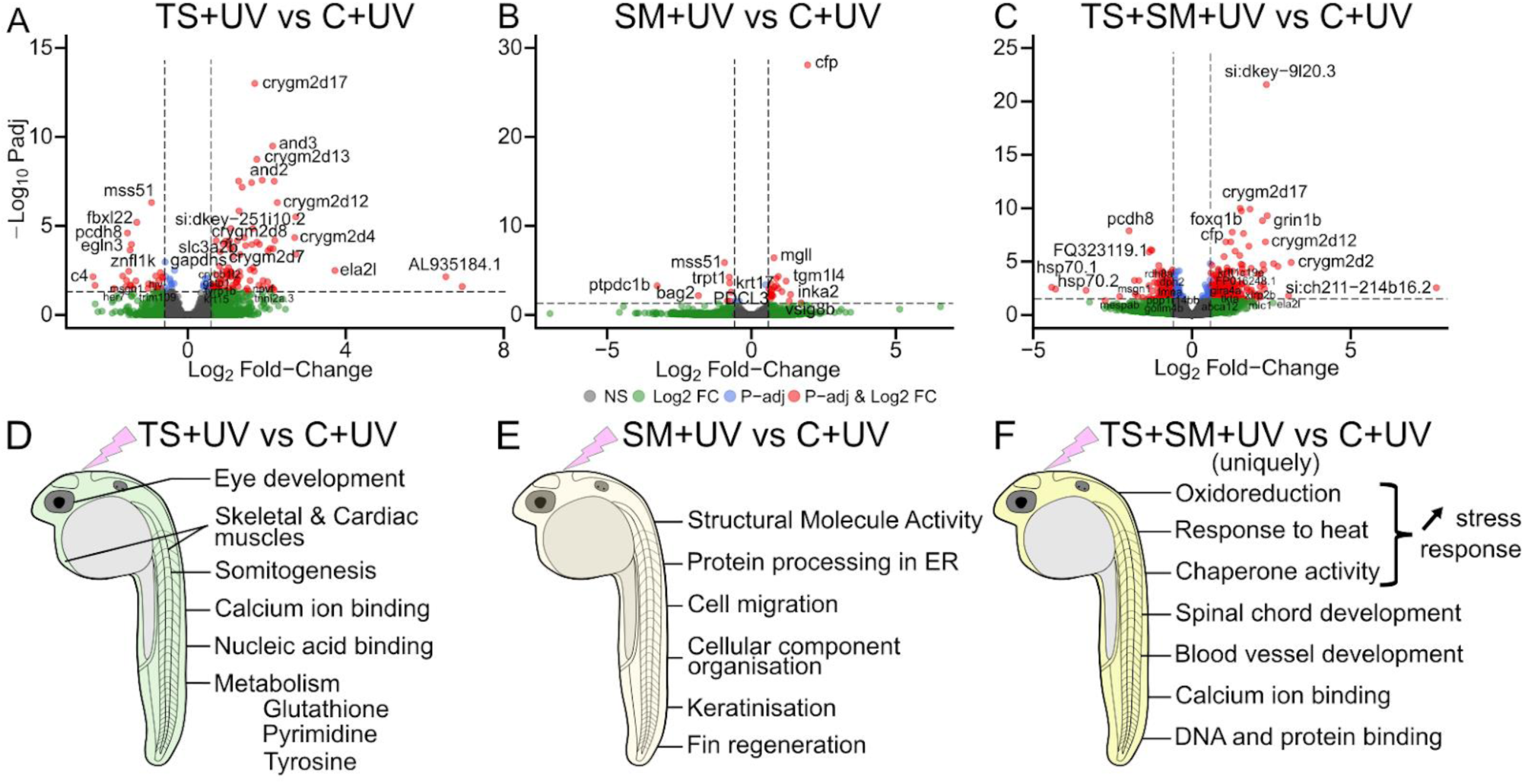
Heat stress and social metabolites followed by UVR alter important molecular pathways. Volcano plots (upper row) showing the differentially expressed genes in response to (A) thermal stress (TS+UV *vs.* C+UV), (C) social metabolites (SM+UV *vs.* C+UV), and (E) their combination (TS+SM+UV *vs.* C+UV). Genes of interest are shown in red when significant (p-adj < 0.05, |FC| > 1.5). Genes left of the left vertical line and right of the right vertical line are underexpressed and overexpressed, respectively, compared to the control C+UV. Functional enrichments (bottom row) of transcriptomic response to TS+UV (B), SM+UV (C), and TS+SM+UV (F) compared to C+UV. Main functionally enriched terms for Gene Ontology of Biological Processes and Molecular Functions, KEGG and Reactome pathways are shown. Treatments were, after UVR exposure, C+UV: control in fresh medium at 27°C, SM+UV: social metabolites at 27°C, TS+UV: fresh medium in thermal stress, TS+SM+UV: social metabolites in thermal stress. Sample size is n = 3 biological replicate pools of 20 embryos per treatment.

Compared to C+UV, heat stress in TS+UV altered 133 genes functionally enriched for eye development, somitogenesis, skeletal and cardiac muscle, macromolecule binding, but also in the metabolism of glutathione (including the glutathione S-transferase *gstp1*), tyrosine (involving *tyrp1a* and *tyrp1b*) and pyrimidine (Figs 3D, S1F). Compared to C+UV, social metabolites in SM+UV induced 31 DEGs involved in fin regeneration, structural molecule activity, and keratinisation, involving several keratin-related transcripts (*krt5/17/92*, *cyt1*, *cyt1l*). Socials metabolites markedly upregulated two immune-related genes, *cfp* and *vsig8b* (Figs 3B,E, S1G). Comparisons within UVR treatments indicated the interacting effect of heat and social metabolites when combined since, compared to C+UV, TS+SM+UV altered the expression significantly more genes (n = 408) than TS+UV (n = 133, χ² = 140.0, p < 0.0001) and SM+UV (n = 31, χ² = 324.0, p < 0.0001, Figs 3C, S1I). Whilst DEGs of TS+SM+UV were partially (n = 94) ascribed to the response to temperature, further 209 DEGs were uniquely found in the combined treatment TS+SM+UV and were involved in temperature response, oxidoreduction, folding activity, macromolecule binding, but also in spinal cord and blood vessel development (Figs 3C,F, S1H,I). Comparing TS+SM+UV to SM+UV and TS+UV provided further evidence for an interactive effect of heat and social metabolites in combination, altering DEGs and pathways distinct from heat or social metabolites alone (Fig. S1J-M).

Stress history altered more genes in response to UVR, with treatment-specific UV-induced DEGs being increased by 60% in SM+UV *vs.* SM (n = 102 genes, χ² = 8.7, p = 0.0032), quadrupled in TS+UV *vs.* TS (n = 264, χ² = 122.0, p < 0.0001), and almost trebled in TS+SM+UV *vs.* TS+SM (n = 171, χ² = 48.7, p < 0.0001), compared to C+UV *vs.* C (n = 64 genes, Fig. 2A,B). The combined treatment TS+SM+UV *vs.* TS+SM induced significantly fewer treatment-specific UV-responsive genes compared to TS+UV *vs.* TS (χ² = 19.9, p < 0.0001), a pattern that held true for the overall number of UVR induced genes (Fig. 2A,B). The 64 genes unique to C *vs.* C+UV, which represent the general response to UVR, were associated with eye, ear, and brain development, hexose metabolism, “response to chemicals” (including peroxiredoxin *prdx1* and another heat shock protein, *hsp90aa1.2*), and nucleotide/carbohydrate binding (Figs 2D, S1N). The 102 genes unique to SM+UV *vs.* SM were enriched for peptidyl-lysine methylation with three notable genes, *eef1akmt2*, *smyd1b*, and *suv39h1a* (Figs 2E, S1O). The 264 genes unique to TS+UV *vs.* TS were associated with gene expression regulation and RNA metabolism, brain and central nervous system development, response to stimulus, which may lead to the metabolism of glutathione (including several glutathione S-transferases *gstt1b, gstp1, gsto2,* and *gstk2*), inositol phosphate, and cholesterol (Figs 2F, S1P). The 171 genes induced by TS+SM+UV *vs.* TS+SM were linked to ribonucleotides, nucleotide di/triphosphates, ion homeostasis, signal transduction, but also with further ribosomal deactivation (with 28 more downregulated rRNAs) and nonsense mediated decay (NMD). Of note, TS+SM+UV also induced several locomotory and swimming behaviour genes, namely *atoh7*, *gdpd5a*, *htr2cl1*, *pleca*, and *pogza* (Figs 2G, S1Q).

### Candidate UV-induced pathways

Stress history-dependent responses to UVR were further explored within candidate pathways associated with macromolecule processing: “RNA integrity Number” (RIN), “DNA repair” (GO:0006281), “protein folding” (chaperone response, GO:0006457), *“*regulation of gene expression, epigenetic*”* (GO:0040029), and “macromolecule methylation” (GO:0043414). UVR degraded RNA, regardless of heat (F = 0.01, p = 0.9192) and medium (F = 0.18, p = 0.6750), as evidenced by lowered RIN values (F = 6.44, p = 0.0195; Fig. S2A). Two DNA repair genes, *eya1* and *fancf*, were respectively down- and up-regulated in all treatments in response to UVR (Fig. 4A,B). Heat stress, but not social metabolites, increased the number (n = 19 *vs.* 7 genes, χ² = 5.54, p = 0.0186) and expression magnitude (H = 4.23, p = 0.0397, “very small effect size” |d| = 0.11) of DNA repair genes, in response to UVR (Fig. 4C).

**Figure 4.**
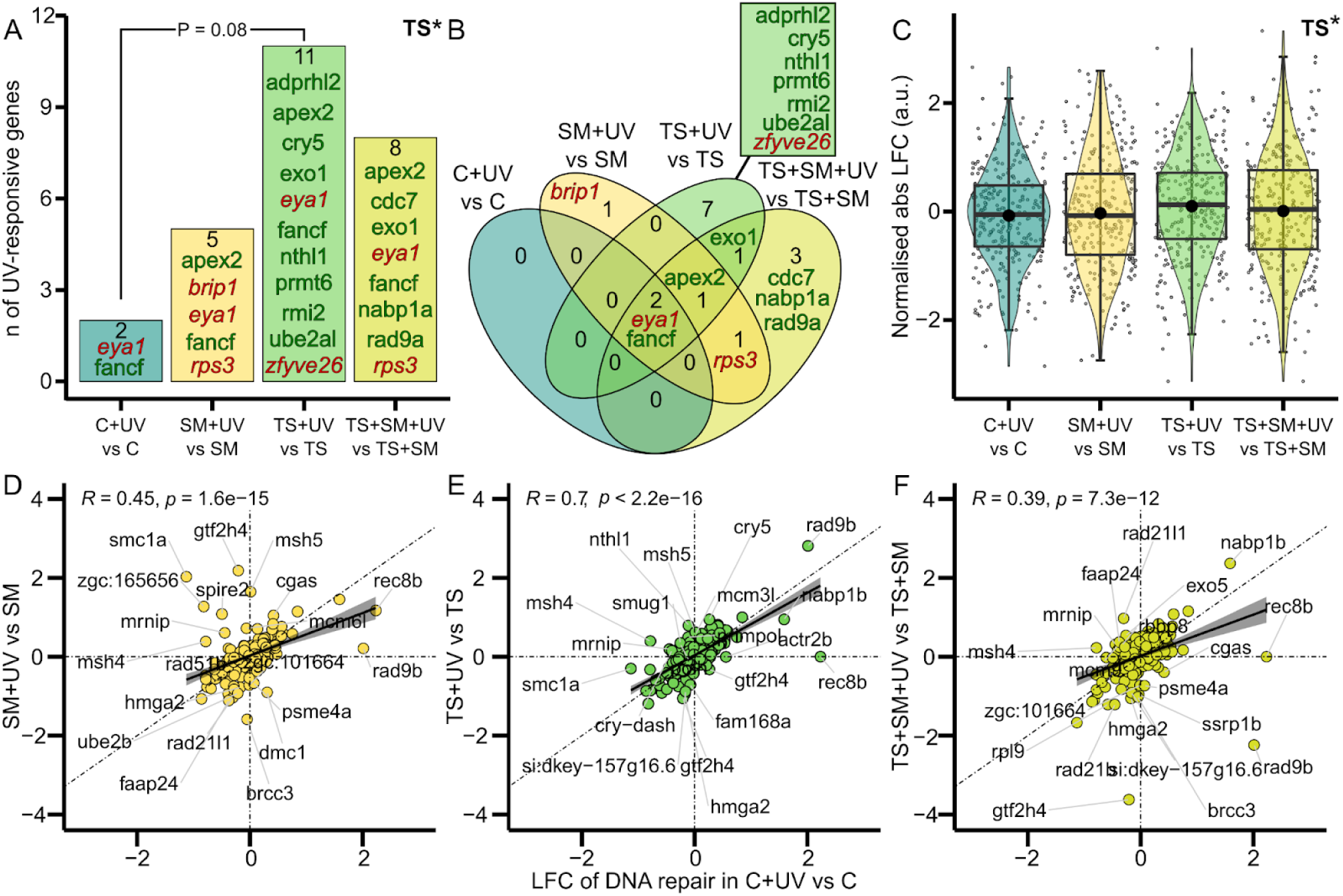
Influence of stress history treatments on DNA repair genes in response to UVR. A) Counts and B) Venn diagram of significant genes from DNA repair (GO:0006281, n = 288 genes) in each treatment in response to UVR. Non-italicised dark green and italicised dark red gene names respectively indicate up- and downregulation (GO-term wide p-adj < 0.05 and |FC| > 1.5). C) Magnitude of UV-induced DNA repair gene expression changes represented by the normalised absolute log-fold change (LFC). Heat stress (TS) is a significant predictor shown on top-right corners in A (Chi-squared test) and C (Scheirer-Ray-Hare test) with * indicating p < 0.05. Bottom row: LFC of gene expression in DNA repair pathway in (D) SM, (E) TS, and (F) TS+SM stress history treatments (y-axes) versus control C (x-axes). The null hypothesis is that the UV-induced repair response is similar for all treatments and respects a 1-to-1 ratio (diagonal lines). Solid black lines and shaded areas depict linear fits and confidence intervals. Labels indicate the top 20 genes with the most pronounced deviation relative to C. Treatments were, before and after (+UV) UVR exposure, C: control in fresh medium at 27°C, SM: social metabolites at 27°C, TS: fresh medium in thermal stress, TS+SM: social metabolites in thermal stress all compared to their non-UV pairs. Sample size is n = 3 biological replicate pools of 20 embryos per treatment.

We expected that heat stress history would limit DNA repair capacities. However, TS+UV *vs.* TS upregulated six (*adprhl2*, *cry5*, *nthl1*, *prmt6*, *rmi2*, and *ube2al*), and downregulated one (*zfyve26*), DNA repair genes not found in C+UV *vs.* C. One DNA repair gene, *apex2*, was significantly upregulated under all stress history conditions but not in C+UV *vs.* C (Fig. 4B). Two DNA repair genes were associated only with social metabolites, with *brip1* downregulated by UV in SM+UV *vs.* SM and *rps3* downregulated in both social metabolites treatments (TS+SM+UV *vs.* TS+SM and SM+UV *vs.* SM but not in TS+UV *vs.* TS; Fig. 4B). Furthermore, deviation from the 1-to-1 expression ratio compared to C+UV *vs.* C evidenced perturbations of the DNA repair pathway by heat stress history (Fig. 4D-F). UV-induced chaperone responses were overall consistent in all treatments both in terms of number of genes and expression magnitude, which were not altered by heat (χ² = 1.47, p = 0.2250 and F = 0.2, p = 0.6736) nor medium (χ² = 0.02, p = 0.8930 and F = 0.0, p = 0.9862, Fig. S2B-G). Seven shared chaperone-related genes included three HSPs (*hsp70l*, *hsp90ab1*, and *hspa8*), but also *dnajb1b, ppiaa, tbcc*, and *zgc:122979* (an hsp 40 family member, a human orthologue of DNAJB5, Fig. S2A-F). The expression of several hallmark genes, such as *hsp70.3, hsp90aa1.2, fkbp1b, dnajb1,* and *dnaja1b,* in TS+SM+UV *vs.* TS+SM was not as strong as the control response to UVR in C+UV *vs.* C; or was already activated by heat stress (without UVR) but not further increased by UVR (Fig. S3F,G). Particularly, *hsp70l* increased by 200-fold in C+UV *vs.* C, but only 6-fold in TS+SM+UV *vs.* TS+SM, representing a 33-fold drop in its expected UVR induction (Fig. S2G). Overall, the proportion and expression magnitude of protein methylation were similar in response to UVR in all four treatments (Fig S2H-J). Nonetheless, social metabolites had an effect on protein methylation as SM+UV *vs.* SM altered transcript levels of three protein methyltransferases (*suv39h1a, eef1akmt2,* and *smyd1b*) not found in C+UV *vs.* C (Fig. S2H,I). Social metabolites (χ² = 5.0, p = 0.0253), but not heat stress (χ² = 0.2, p = 0.6550), activated several epigenetic regulators, with four genes found in social metabolites treatments: *bmi1b*, *srrt*, and two histone linkers, *h1-0* and *si:ch73-368j24.12* (human ortholog: H1.5 linker histone; Fig. S2N). Social metabolites also altered the expected 1–to-1 expression ratios of epigenetic genes between SM+UV *vs.* SM and C+UV *vs.* C (R = 0.18, p = 0.098, Fig. S2P). Of note, heat marginally tended (weak evidence: p < 0.1) to increase the magnitude of gene expression of both methylation and epigenetic processes (Figs S2J,O).

### Heat rescued phenotypes, whilst social metabolites caused hypoactivity and TS+SM overwhelmed larvae

There were no differences in mortalities, with > 95% of embryos surviving the UVR exposure regardless of treatments (Fig. S3A). We tested the hypothesis that heat stress history causes more mutant phenotypes in response to UVR through measuring fitness-relevant outcomes (Figs 5A-E,H; all statistical analyses of phenotypic data are reported in Dataset S1).

**Figure 5.**
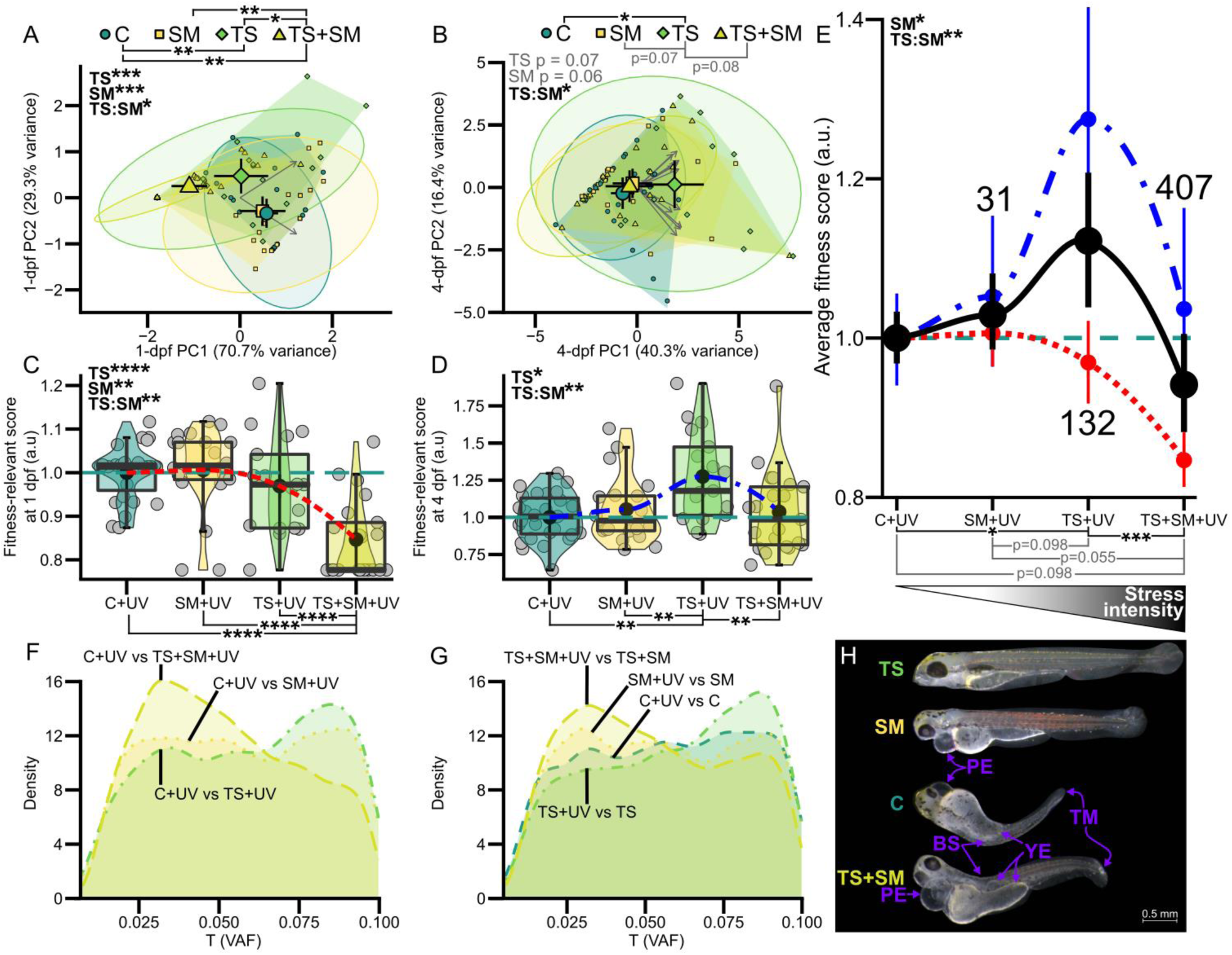
TS+UV outperformed C+UV and TS+SM+UV showing a nonlinear response to ultraviolet radiation (UVR) with stress intensity. Principal Component Analyses of phenotypes at 1 (A) and 4 (B) days post fertilisation (dpf). Higher values along PC1 indicate higher fitness-relevant outcomes. Large and small coloured symbols show centroids (± 95% confidence interval, CI) and individual scores, respectively. Ellipsoids show 95% normal CI probabilities. Polygons show smallest areas clustering samples per treatment. Fitness-relevant scores (PC1 scores) are shown in function of increased stress intensityat 1 (C) and 4 (D) dpf. Boxes show median and 25%-75% quartiles. Whiskers are min/max values within 1.5 interquartile range. Individual data given as jittered grey circles. (E) Average fitness-relevant scores (means ± s.e.m) from 1 and 4 dpf. Smooth *loess* lines show 1-dpf (dotted red line), 4-dpf (dot-dashed blue line), and average (black solid line) fitness-relevant scores. Black dots show mean ± s.e.m in arbitrary units (a.u.) scaled to control (C+UV, dashed blue horizontal lines) for neatness. Numbers in (E) show how many genes were altered compared to C+UV. Significant 2-way model terms (PERMANOVAs in A-B and ANOVAs in (C-E) are shown in top-left corners with TS = thermal stress and SM = social metabolites. Post-hoc comparisons are shown with horizontal bars. Density plots of SNVs (variant allele frequency) comparing (F) treatments with their non-UV treated counterparts, and (G) UV-assayed treatments with UVR-exposed controls. In each comparison group, TS+SM had the highest number of SNVs whilst TS had the lowest number. Sample size is n = 3 biological replicate pools of 20 embryos per treatment. (H) Compared to C, embryos have fewer defects and are longer in TS but show larger pericardial-to-embryo length ratios with social metabolites SM and TS+SM. Defects include bent spine (BS), pericardial edema (PE), tail malformation (TM), and yolk extension disruption (YE). Image modification: selected embryo images were manually detoured, contrast adjusted evenly for all treatments, and put on a black background. All tests are two-tailed with adjustment for False Discovery Rate when multiple comparisons are made. Treatments were all followed by UVR (+UV), with C+UV: control in fresh medium at 27°C (n = 28), SM+UV: social metabolites at 27°C (n = 22), TS+UV: fresh medium in thermal stress (n = 20), TS+SM+UV: social metabolites in thermal stress (n = 25). *: p ≤ 0.05, **: p ≤ 0.01, ***: p ≤ 0.001, ****: p ≤ 0.0001. Weak evidence (p < 0.1) is shown in grey. Phenotypic analyses were replicated once in the laboratory.

The 1-dpf multivariate data showed that both heat (F = 28.0361, p = 0.001) and social metabolites (F = 9.14, p = 0.001) altered 1-dpf startle behaviours, with significant differences between TS+UV (F = 7.78, p = 0.006) and TS+SM+UV (F = 34.74, p = 0.006) compared to C+UV (Fig. 5A). 1-dpf embryos in TS+SM+UV behaved differently from SM+UV (F = 17.99, p = 0.006) and TS+UV (F = 8.27, p = 0.024). Conversely, the 4-dpf multivariate data showed that only TS+UV differed from C+UV (F = 5.62, p = 0.03), with weak evidence that TS+UV also deviated from SM+UV (F = 4.72, p = 0.078) and TS+SM+UV (F = 4.69, p = 0.066, Fig. 5B). We then individually analysed phenotypic variables at 1 and 4 dpf (Fig. S3A-P). Both heat stress (H = 38.1, p < 0.0001) and social metabolites (H = 3.96, p = 0.0465) reduced light responsiveness in 1-dpf embryos (Fig. S3E,F), with TS+UV (W = 457, p = 0.0009) and TS+SM+UV (W = 661, p < 0.0001) lowering burst activity % compared to C+UV (Fig. S3E). Larvae in SM+UV (t = −2.3, p = 0.0491) and TS+UV (t = 3.8, p = 0.0016) were longer relative to C+UV (Fig. S3G). In turn, heat stress (z = 2.8, p = 0.0051) facilitated hatching, with an increase from 11% in C+UV to 50% in TS+UV (z = −2.8, p = 0.0306, Fig. S3B). There was weak to moderate evidence that social metabolites (F = 4.9, p = 0.0287) and heat (F = 2.9, p = 0.0945) altered the 4-dpf embryo size-normalised pericardial width (Fig. S3J). Larvae in TS+UV had narrower pericardia relative to their body sizes compared to C+UV (t = 2.91, p = 0.0104) and SM+UV (t = 2.87, p = 0.0104). There was strong evidence for protective effects against pericardial edema with heat stress (z = −3.3, p = 0.0010), with a drastic decline in pericardial edemas only in TS+UV (32%, z = 3.3, p = 0.0059) compared to C+UV (82%, Fig. S3C) — although heart rates remained similar across treatments (Dataset S1). Larvae exposed to TS+UV also had the lowest defect score, albeit not significant relative to C+UV (Fig. S3I). Heat stress (F = 8.27, p = 0.0051) also significantly increased the eye-to-body size ratio following exposure to UVR (Fig S3H). At 4 dpf, there was weak to moderate evidence that social metabolites reduced the dark-light swimming activity (H = 3.86, p = 0.04955, Fig. 3K) and swimming burst counts (H = 3.37, p = 0.0663, Fig. S3M) and touch-evoked swimming speed (F = 3.64, p = 0.0599, Fig. S3P). Conversely, despite having experienced UV, 4-dpf larvae in TS+UV were constantly most responsive and had the highest number of active larvae, activity %, distance swam, burst count, and speed in response to light (Fig. S3D,K-N); but also, the highest distance swam and speed in response to touch (Fig. S3O,P). Interestingly, the morphological consequences of exposure to TS+UV were associated with higher swimming responsiveness with positive (embryo size) and negative (pericardial edema, pericardial width, and defect scores) relationships. Conversely, embryos in the combined treatment TS+SM+UV lost the developmental effect conferred by heat and social metabolite treatments alone and were smaller compared to TS+UV (t = 3.3, p = 0.0035) and SM+UV (t = 1.89, weak evidence: p = 0.0937), reaching similar sizes as in the control C+UV (t = −0.32, p = 0.7474, Fig. S3G). The heat-induced protection against wide pericardia was lost in TS+SM+UV compared to TS+UV (t = −3.11, p = 0.0104), resulting in similar pericardial width relative to C+UV (t = −0.35, p = 0.8889, Fig. S3J). This caused twice as much pericardial edema (although with weak evidence, z = −2.12, p = 0.1008, Fig. S3C) in conjunction with higher defect scores (W = 116, p = 0.0490, Fig. S3I) in TS+SM+UV compared to TS+UV. Likewise, TS+SM+UV caused the lowest responsiveness to light at 1 dpf (Fig. S3E-F) and to light and touch stimuli at 4 dpf (Fig. S3K-P). Altogether, this indicated that the combined treatment TS+SM+UV induced more defects and negative outcomes compared to TS.

Summarising the multivariate data into a “fitness-relevant score” (where higher values indicate better fitness-relevant outcomes) confirmed that TS+SM+UV performed worst compared to C+UV (t = 5.92, p < 0.0001), SM+UV (t = 5.72, p < 0.0001), and TS+UV (t = 4.22, p < 0.0001) and displayed the lowest fitness-relevant scores at 1 dpf (Fig. 5C). On the other hand, TS+UV increased fitness-relevant scores at 4 dpf compared to C+UV (t = −3.47, p = 341 0.0049), SM+UV (t = −2.54, p = 0.0257), and TS+SM+UV (t = 3.05, p = 0.0094, Fig 5D). Overall, the average fitness-relevant scores peaked with TS+UV, which had the highest average fitness-relevant score, higher than C+UV (t = −2.51, p = 0.0422) and TS+SM+UV (t = 3.9, p = 0.0010). The average fitness-relevant scores dropped in the multistress treatment TS+SM+UV, which tended to have lower scores compared to C+UV (t = 1.76, weak evidence: p = 0.0983) and SM+UV (t = 2.24, weak evidence: p = 0.0550, Fig. 5E). In summary, (i) larvae in SM+UV grew longer but were hypoactive at 4 dpf, (ii) larvae in TS+UV showed less pericardial edema and developed faster, which facilitated their hatching and improved their behavioural responses at 4 dpf, (iii) whilst embryos and larvae in TS+SM+UV failed to limit the formation of pericardial edema and had the worst fitness-relevant performances. This indicated a nonlinear relationship between fitness-relevant performances in response to UVR and the intensity of stress history (Fig. 5E).

### Minor Allele Frequencies differed depending on heat stress history

Minor Allele Frequency distributions of treatments in the 0.025 range clearly differed between treatments (Fig. 5F-G, Dataset S1). Comparing UV-exposed *vs.* their non-UV-treated embryos showed that the TS+UV treatment had the lowest MAF at 0.025, followed by the control condition C+UV. Both SM+UV and TS+SM+UV treatments had pronounced MAF peaks at 0.025, indicating spontaneous DNA mutations at the single nucleotide level. All distributions significantly differed from each other (Fig. 5F). In the comparison of each UVR treatment against the control C+UV, the TS+UV treatment again had the lowest MAF at 0.025, with a marginally higher MAF for C+UV against SM+UV, and the highest MAF again observed in the C+UV against TS+SM+UV treatment. Kolmogorov–Smirnov tests showed that the distribution of MAF was significantly different between all four treatments in response to UVR (Fig. 5G, Dataset S1). Treatment-specific UV-induced mutations resulted in a higher number of normal phenotypes in TS+UV whilst defect scores significantly tripled in larvae exposed to TS+SM+UV (W = 116.5, p = 0.049, Fig. S3I), resulting in a higher number of mutant phenotypes with pericardium, yolk, spine, and tail malformations (Fig. 5H).

## Discussion

We explored the effects of stress history on the UVA-catalysed recovery from UVB-induced damage in zebrafish embryos. UVR is known to cause oxidative stress toxic to lipids, antioxidants, ribosomes, proteins, and nucleic acids (Dahms and Lee, 2010; Hurem et al., 2018; Iordanov et al., 1998). Here, UVR altered the metabolism of macromolecules (ribosomes, mRNA, protein), DNA checkpoints, and organ development. The unfolded protein response and ubiquitination pathways protect proteins and remove damaged ones (Bianchi et al., 2015; Simon et al., 1995), likely explaining their activation by UVR in our study. UVR lowered RNA integrity and activated nonsense-mediated mRNA decay and DNA checkpoints, indicating that UVR damaged nucleic acids. The DNA repair response involved *fancf* and *eya1* in all treatments*. fancf* helps recruiting DNA repair genes upon DNA damage (Grompe and D’Andrea, 2001; Kowal et al., 2007; Yao et al., 2015). However, upregulation of only one DNA repair gene, *fancf,* suggests a minimal repairing capacity in control embryos. Moreover, *eya1* is a DNA repair-promoting UVR-inducible enzyme that was downregulated by UVR, indicative of genotoxicity (Cook et al., 2009; Farrell et al., 2011; Zhou et al., 2017). Many cell components were altered by UVR, leading to the activation of autophagy pathways to remove damaged macromolecules (Sample and He, 2017). These cellular effects escalated to the organismal level, resulting in hypoactivity, teratogenicity, and reduced hatching, as previously reported in zebrafish embryos (Charron et al., 2000; Hurem et al., 2018).

Direct heat stress changed this UVR damage/repair response compared to UVR-exposed control embryos. Heat-stressed larvae were longer with fewer pericardial edema, better fitness-related swimming performances, and earlier hatching, providing evidence that heat rescued and/or protected embryos from UVR. These results do not support our initial hypothesis that heat stress would disadvantage embryos in a mutagenic environment. Our results align with concepts of “hormesis” or “cross-protection”, that refer to beneficial stimulating effects of low-dose often mild initial stress (Calabrese and Baldwin, 2002; Costantini et al., 2010; Rodgers and Gomez Isaza, 2023). In our study, the heat hormetic effect rescuing embryos from UVR may have resulted from heat-induced defence mechanisms protecting against and/or repairing irradiation damage. Comparing transcriptomes of heat-stressed embryos before and after the UVR damage/repair assay suggests that hormesis may have been mediated via (i) a stage-dependent increased baseline tolerance, and by stimulating (ii) antioxidants, (iii) the heat shock response (HSR), and (iv) DNA repair.

Heat-stressed embryos were six hours older than control embryos that were still undergoing segmentation when UVR started (Feugere et al., 2023). Older heat-stressed embryos may display different phenotypes from stage-matched controls (Feugere et al., 2021b), but these additional controls could not be added to this experiment, which warrants future research to disentangle the effects of heat and development in the observed response to UVR. Mature fish are more resistant to UVR than earlier stages, possibly through a protective role of pigmentation (Alves and Agustí, 2020; Charron et al., 2000; Dahms and Lee, 2010). Heat treatment upregulated two genes (*tyrp1a/b*) involved in melanin synthesis (Braasch et al., 2009; Krauss et al., 2014), which may shield cells from UVR (Brenner and Hearing, 2008). Overdeveloped heat-treated embryos may also have benefited from energy supply from the yolk sac (Sant and Timme-Laragy, 2018). Likewise, heat upregulated several genes involved in glycolysis (e.g., *gapdh*, *gpia*, *pkmb*) and energy metabolism (Feugere et al., 2023). Therefore, heat-stressed embryos may mobilise energy to fuel ATP-dependent DNA repair mechanisms such as nucleotide excision repair (Reef et al., 2009) and mismatch repair (Groothuizen and Sixma, 2016; Jiricny, 2013). Supporting this, heat-stressed embryos were hypoactive immediately after UVR, suggesting that they may have redirected energy towards UVR-coping strategies.

Antioxidants may mediate hormesis through “preparation for oxidative stress”, an adaptive physiological mechanism that upregulates antioxidants to confer tolerance against stress-induced reactive oxygen species (Costantini et al., 2012; Giraud-Billoud et al., 2019; Oliveira et al., 2018). Supporting this, heat stress upregulated several glutathione-S-transferases (GSTs) following UVR. GSTs play a key role in protecting cellular macromolecules by catalysing the binding of reduced glutathione to damaged macromolecules (lipids, DNA, and proteins), marking them for degradation (Griffiths et al., 1998; Hayes and McLellan, 1999).

The HSR may mediate hormesis by limiting molecular damage (Jantschitsch and Trautinger, 2003; Rattan, 2006). Our heat stress protocol activated the HSR at 1 dpf (Feugere et al., 2021b; 2023), which may have provided heat-treated embryos with a reserve of HSP transcripts ready to be translated once UVR started. HSPs are also key modulators of DNA repair genes, which may in turn favour cell survival (Sottile and Nadin, 2018). Furthermore, heat activated the ubiquitin-proteasome system (UPS, involving several ubiquitins, e.g. *ube2al*, *otulina*, *igs15*, and *otud7b* genes), which plays a central role in DNA repair (Sakai et al., 2020). Therefore, the cellular stress response activated by both heat and UVR may have acted as a short-lived “transient cross-protection” resulting in the cross-tolerance against UVR (Rodgers and Gomez Isaza, 2023).

Unexpectedly, we found that heat stress history improved DNA repair capacity in response to UVR. These findings agree with increased DNA repair rates at high temperature in daphnia (MacFadyen et al., 2004), tadpole (Morison et al., 2020), and zebrafish embryos (Chien et al., 2020). DNA damage not repaired within 2 hours is irreversible in zebrafish embryos (Dong et al., 2008), suggesting that transcriptomic changes captured approximately 30 min following UVR played a key role in mediating hormesis. Whilst control embryos activated one DNA repair gene (*fancf*), heat upregulated several DNA repair genes. One such gene that does not support our initial hypothesis is the evidence for a more competent heat-induced photorepair indicated by the upregulation of *cry5*, a 6-4 photolyase, that may have acted against UVR damage at the time of (UVA and blue) light exposure in TS+UV and may in turn cause higher survival in response to UVR (Banaś et al., 2020; Hirayama et al., 2009; Steindal and Whitmore, 2020; Tamai et al., 2004; Weger et al., 2011). Heat also upregulated other DNA repair-related genes that may have helped embryos in TS+UV to repair DNA damage: *adprhl2, apex2, exo1, nthl1, rmi2,* and *ube2al* (Table 1).

**Table 1.** Heat stress facilitates DNA repair, rescuing embryos from UVR-induced DNA damage.

| Gene | Role | Reference |
| --- | --- | --- |
| *adprhl2* | Prevents cell death by removing poly-ADP ribose accumulated following oxidative stress | Ghosh et al. (2018) |
| *apex2* | Limits DNA damage by preventing the stabilisation of UV-induced DNA cleavage | Lanza et al. (1996); Subramanian et al. (1998); Zaksauskaite et al. (2021) |
| *exo1* | Mediates a faster DNA mismatch repair pathway | Goellner et al. (2015) |
| *nthl1* | Cleaves oxidative pyrimidine damage during base excision repair | Das et al. (2020) |
| *prmt6* | Methyltransferase preventing apoptosis and rescuing embryogenic defects, and regulating DNA polymerase and DNA repair genes | Chen et al. (2022); El-Andaloussi et al. (2006); Zhao et al. (2016) |
| *rmi2* | Promotes genome stability within the Bloom syndrome (BLM) complex, involved in homologous repair | Hudson et al. (2016); Patel et al. (2017); Xu et al. (2008) |
| *ube2al* | E2 ubiquitin-conjugating enzymes such as *ube2a* mediate protein ubiquitination promoting DNA repair and genome integrity | Brinkmann et al. 2015; Cukras et al. (2014); Stewart et al. (2016) |

Likewise, heat stress activated the metabolism of inositol phosphate, which is known to stimulate double strand break repair through non-homologous end joining (Hanakahi et al., 2000). Furthermore, heat stress increased (but with weak evidence) the expression of transcripts involved in epigenetics and methylation. Histone methylation can transmit heat hormetic effects to later life stages and subsequent generations (Wan et al., 2021), which may mean that heat-treated embryos not only perform better in response to UVR, but could also be better protected from subsequent stress events.

Overall, heat stress history rescued embryos from molecular damage in a mutagenic environment, likely by chaperoning proteins, activating DNA repair, stabilising the genome, and facilitating methylation and epigenetic regulation. This was supported by fewer SNVs in heat-treated embryos compared to all other treatments. Whilst our findings align with the concept of hormesis wherein heat protects cells by activating the proteasome, the HSR, and antioxidants (Haarmann-Stemmann et al., 2013; Rattan, 2006), further research is needed to discriminate the involvement of protective *vs*. repair pathways in the hormetic effect observed here. We also acknowledge limitations to using behaviour and morphology as fitness proxies (Cocciardi et al., 2021; Higginson, 2020; Lind and Cresswell, 2005) and that future studies will need to assess whether these fitness advantages hold in the natural environment, particularly if hormesis is mostly mediated by the transient cellular stress response, as discussed in Rodgers and Gomez Isaza (2023). The loss of thermal plasticity and UVB tolerance in laboratory-adapted zebrafish strains (Dong et al., 2007; Morgan et al., 2022) and the low percentage of explained variance (40.3%) in the 4-dpf phenotypic data also warrants repeated experiments validating our findings using wild zebrafish lines. Of note, severe heat exacerbates stress-induced damage, following an inverted U-shaped dose-response curve (Calabrese and Baldwin, 2002; Haarmann-Stemmann et al., 2013; Matsuda et al., 2013). The positive outcome of heat stress observed here is likely explained by the sublethal heat regime we used. We hypothesise that temperatures beyond 32°C would instead lead to negative outcomes in a mutagenic environment, which warrants further investigation.

Biotically-stressed aquatic animals communicate with naive neighbours (Crane et al., 2022; Mathuru, 2016; v. Frisch, 1938). We previously extended this mechanism to abiotic stress and identified specific classes of social metabolites (Feugere et al., 2021a; 2021b; 2023). Here, we confirm that heat stress responses can be indirectly transmitted between embryos via social metabolites activating immune-, cell structure- and keratin-related genes in receivers. Similar to direct heat stress, embryos in SM+UV were hypoactive at 1 dpf and longer at 4 dpf, but the activated molecular pathways were markedly different from those activated by heat stress. Stress history conditions, including social metabolites, but not control conditions, activated *apex2*, an important DNA repair gene in embryogenesis (Fortier et al., 2009; Zaksauskaite et al., 2021). Keratin genes activated by social metabolites may have improved photoprotection against DNA damage in keratinocytes (Wondrak et al., 2006; Wu and Hammer, 2014) and promoted repair of DNA breaks (Nair et al., 2021). However, the response to UVR in the social metabolites treatment also downregulated both *brip1* and *rps3*, suggesting some level of DNA damage (Bridge et al., 2005; Elsakrmy et al., 2022) that may have caused the observed malformations. Social metabolites altered epigenetic and methylation genes, which may mediate downstream transcriptomic changes (Gibney and Nolan, 2010). For instance, SM+UV downregulated *srrt*, which plays a role in DNA repair (Bui et al., 2019; Rossman and Wang, 1999), histone h1-5, and methyltransferases, suggesting alterations in transcriptional regulation (Albig et al., 1997; Vougiouklakis et al., 2020). One differentially expressed gene was *smyd1b*, playing a key role in skeletal muscles (Li et al., 2013; Tan et al., 2006), the downregulation of which may explain how social metabolites rendered embryos hypoactive. Therefore, while social metabolites induced markedly more responsiveness to UVR compared to controls, just like heat stress did, they failed to protect embryos against UVR damage as efficiently as heat stress. While the effects observed in the social metabolite treatment could be ascribed to specific metabolites induced by heat, similar to what we reported previously ((Feugere et al. 2021; Feugere et al. 2023)) we cannot exclude that this bouquet of cues (Feugere et al. 2023) also includes regularly excreted, non-heat-specific social cues. Ideally, cues from unstressed conspecifics should be included in future experiments as a better-suited control (Crane et al., 2022) to confirm the involvement and consequences of heat-induced vs. regular social metabolites in mutagenic environments.

The TS+SM treatment combining heat stress within a social context more profoundly altered the transcriptome both before (Feugere et al., 2023) and after UVR exposure. One possible explanation for this is that single exposure to either direct heat stress (TS) or the social context of heat stress (social metabolites) influenced distinct pathways in response to UVR, which would both be activated in the combined treatment. These embryos experienced increased stress levels as suggested by more stress-related functional changes, further damage to RNA, and more inhibition of translation by downregulating ribosomal RNAs. The combined treatment elevated HSPs levels before UVR (Feugere et al., 2023). Such elevated basal HSP mRNAs may have attenuated the HSR to subsequent acute stressors (Sessions et al., 2021; Whitehouse et al., 2017). Supporting this idea, the combined treatment limited and/or inhibited the upregulation of key HSPs in response to UVR, likely lowering protection against protein damage. The interacting effect of heat and social metabolites amplifying transcriptome changes also likely cost more energy, as suggested by more pronounced behavioural hypoactivity. Overall, our data suggest that the combination of direct heat with social metabolites was a stronger stressor that prevented embryos from activating the heat hormetic effect, and shifted the dose-response towards negative outcome. This was supported at the sequence level with this treatment having the most SNVs compared to UV-treated controls and compared to TS+SM treatments without UVR. It may follow that in a natural environment, where embryos in a clutch exchange social metabolites, any protective hormetic effect of heat will be negated by the presence of social metabolites exchanged between clutch mates. Our results may suggest that species that are found or raised in high densities would be disadvantaged in stressful environments if they had experienced both direct stressors and social metabolites in early life stages.

## Conclusion

Our contribution evidenced a nonlinear stress/fitness relationship characterised by positive (body size) and negative (defects) outcomes in chemically-communicated indirect heat stress, by a protective hormetic effect activated by direct sublethal heat stress, and by negative fitness-relevant outcomes in embryos experiencing combined stress. Our finding that stress history can alter the response to UVR is relevant for natural populations, which must navigate increasingly common heat events in mutagenic environments that may be exacerbated by climate change (Bais et al., 2018; Ficke et al., 2007). For instance, the fact that sublethal heat stress may increase the tolerance to UVR warrants further research as cross-protection may facilitate “pre-adaptation” for organisms facing increasing anthropogenic stressors (Rodgers and Gomez Isaza, 2023). In conclusion, we showed that the response to a mutagenic environment depended on the type and intensity of heat stress experienced during early development.

## Acknowledgements

We thank the technical staff from the Biology and Marine Sciences department at the University of Hull for providing us with the equipment for experimental ultraviolet exposure. We would like to thank Alan Smith for fish care in the aquarium facility at the University of Hull. We acknowledge the Viper High Performance Computing facility of the University of Hull and its support team, in particular Chris Collins and Darren L. Bird.

## Author contribution

KWV designed the study. LF acquired experimental data. LF, AB, CSDF, and KWV analysed the data. LF wrote the first draft and all authors incl. PBA and KBS contributed to the final version.

## Data availability

The data for phenotypic experiments, processed transcriptomic data, and custom code (R scripts and bioinformatic pipelines) will be made available upon publication on Zenodo (reserved accession number: 10.5281/zenodo.7566285). The raw sequences from RNA-sequencing will be made available on NCBI Gene Expression Omnibus under the accession number GSE223685 upon publication. The raw transcriptomic data of the embryos not exposed to UVR is available on NCBI Gene Expression Omnibus under the BioProject PRJNA910181 and the accession number GSE220546.

## Ethics statement

All experiments were approved by the Ethical committee of the University of Hull under the Ethics reference FEC_2019_194 Amendment 1.

## List of Symbols and Abbreviations

ANOVA: analysis of variance.
ATP: adenosine triphosphate
C: control
CNS: central nervous system
CPDs: cyclobutene pyrimidine dimers
DEGs: differentially expressed genes
dpf: day post fertilisation
FC: fold change
GST: glutathione S-transferase
hpf: hour post fertilisation
HSP: heat shock protein
HSR: heat shock response
LFC: log fold change
NMD: non-sense mediated decay
PC: principal component
PCA: principal component analysis
PERMANOVA: permutational multivariate analysis of variance
RIN: RNA integrity number
s.e.m.: standard error of the mean
SM: social metabolites
SNVs: single-nucleotide variants
TS: thermal stress
UV: ultraviolet
UVA: ultraviolet A
UVB: ultraviolet B
UVR: ultraviolet radiation
6-4PPs: 6-4 photoproducts
|d|: effect size

