## Supplementary Information for "Consequences of directly- and indirectly-experienced heat stress in a mutagenic environment"

### Supplementary Methods

We aimed to optimise the ultraviolet B (UVB) and ultraviolet A (UVA) setup (UVB/UVA) to induce sufficient DNA damage/repair whilst avoiding substantial mortality up to 5 days. Several optimisation steps were necessary before an acceptable UVB/UVA setup was obtained. We optimised an in-house setup to best match the results of Dong and collaborators (Dong et al., 2008) who showed complete recovery of UVB induced alterations by UVA photorepair. For this purpose, non-stressed control zebrafish embryos were exposed to UVB only (for 3, 6, 9 min with a single irradiation source), UVA only (28 min with a single irradiation source), and UVB+UVA. The best UVB exposure time was determined to be 6 min as that of 9 min induced too much death and strong defects that would not be recovered. Preliminary experiments showed that UVB induces mortality, mutations and defects on phenotype. These deformities are: heart oedema, no caudal fin formation, development delay, yolk ball and extension alteration, and lateral fin alteration. These defects are less severe, delayed or recovered when subjected to subsequent UVA at 1 dpf. Such UVB+A treatment allowed complete avoidance of mortality up to 4 dpf. There were no defects caused by UVA only.

In order to catch transcriptomic changes occurring just after UVA repair, there should not be too much delay after the UV treatments. Hence, the 28 min of bottom-up UVA exposure was replaced by 15 min of bottom-up/top-down UVA exposure with a double source. A small set of preliminary data showed that UVA allows recovery from otherwise UVB-induced mortality, although there were still defects that were not completely recovered as shown by the presence of heart oedema and curvature in UVB+UVA treated embryos. The successful photorepair effect of UVA was confirmed with a significant increase in the ratio of healthy embryos in UVB+UVA (n = 14/21, 61%), to similar levels as in the control (n = 19/21, 90%, $\chi^2$ = 0.76, p-adj = 0.3840), compared to UVB (n = 0/24, 0%, $\chi^2$ = 14, p-adj= 0.0004), see "Statistics", Dataset S1). Embryos exposed to UVB+UVA treatment (n = 23/23, 100%) increased their survival, to similar levels as in the control (n = 21/21, 100%, $\chi^2$ = 0.1, p-adj = 0.763), compared to embryos exposed to UVB alone (n = 14/24, 58%, $\chi^2$ = 5.12, raw p = 0.0236, p-adj = 0.0708, see "Statistics", Dataset S1).

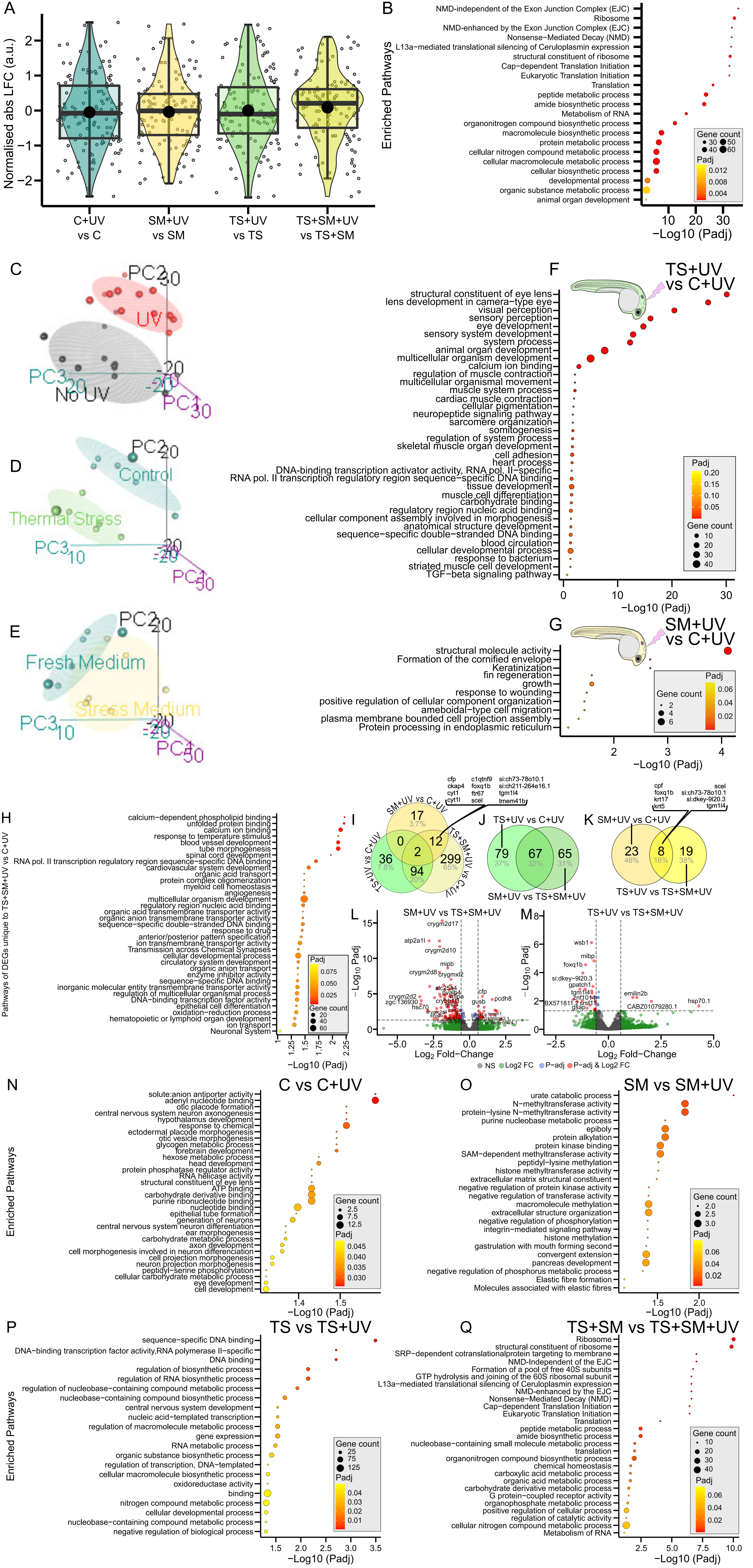


**Figure S1. The transcriptomic response to UVR has common and unique components depending on heat stress history.**
(S1A) Box-Cox normalised absolute log-fold change (LFC) values of shared genes activated by ultraviolet radiation (UVR) in function of treatments. Black dots show mean ± s.e.m in arbitrary units (a.u.). Boxes show median and 25%-75% quartiles. Whiskers are min/max values within 1.5 interquartile range. Individual genes are given as jittered grey circles. (S1B) Functional enrichment showing top functional terms associated with differentially expressed genes (DEGs) shared by all UVR treatments (C+UV, SM+UV, TS+UV, TS+SM+UV) compared to their non-UV pairs. (S1C-E) Principal component analyses (PCA) showing the first three components of the spectral decomposition of the regularised log transformed variance of the read count data. PCAs compare (S1C) UVR (red ellipse and dots, n = 12) vs. no-UVR (black ellipse and dots, n = 12) samples;(S1D) thermal stress (in green, n = 6) vs. control temperature (in blue, n = 6) within UVR treatment; and (S1E) social metabolites (in yellow, n = 6) vs. fresh medium (in blue, n = 6) within UVR treatment. (S1F-G) Functional enrichments showing top functional terms of the DEGs found in (S1F) TS+UV and (S1G) SM+UV compared to C+UV.(S1H) Functional enrichment showing top functional terms of the 299 DEGs found in TS+SM+UV but not in SM+UV and TS+UV. (S1I) Venn diagram of significant DEGs shared vs unique to treatments compared to C+UV with many genes unique to TS+SM+UV. (S1J) Venn diagram of DEGs associated with thermal stress and shared/unique between [TS+UV vs C+UV] and [SM+UV vs TS+SM+UV] (i.e. effect of TS). (S1K) Venn diagram of DEGs associated with social metabolites and shared/unique between [SM+UV vs C+UV] and [TS+UV vs TS+SM+UV] (i.e. effect of SM). (S1L-M) Volcano plots of (S1L) SM +UV vs TS+SM+UV and (S1M) TS+UV vs TS+SM+UV evidencing an interactive effect of heat and social metabolites on the transcriptomic response to UVR. Volcano plots show DEGs in red when significant (p-adj < 0.05, |FC| > 1.5). DEGs left to the left vertical line and right to the right vertical line are respectively significantly under- and over-expressed compared to TS+SM+UV. (S1N-Q) Stress history induced unique transcriptomic responses to UVR evidenced by functional enrichment showing top functional terms of DEGs unique to each UVR treatments compared to their non-UV pairs in (S1N) C vs C+UV, (S1O) SM vs SM+UV, (S1P) TS vs TS+UV, and (S1Q) TS+SM vs TS+SM+UV. Functional enrichments represent terms based on highest gene count/term with n ≥ 2 genes/term, and significant DEGs with p-adj < 0.05 and |FC| > 1.5. Gene Ontology terms for Biological Processes and Molecular Functions, as well as KEGG pathways and Reactome pathways are represented. Treatments were C: control in fresh medium at 27°C, SM: social metabolites at 27°C, TS: fresh medium in thermal stress, TS+SM: social metabolites in thermal stress, all compared to their non-UV pairs. EJC: exon junction complex. Pol.: polymerase. SAM: S−adenosylmethionine.

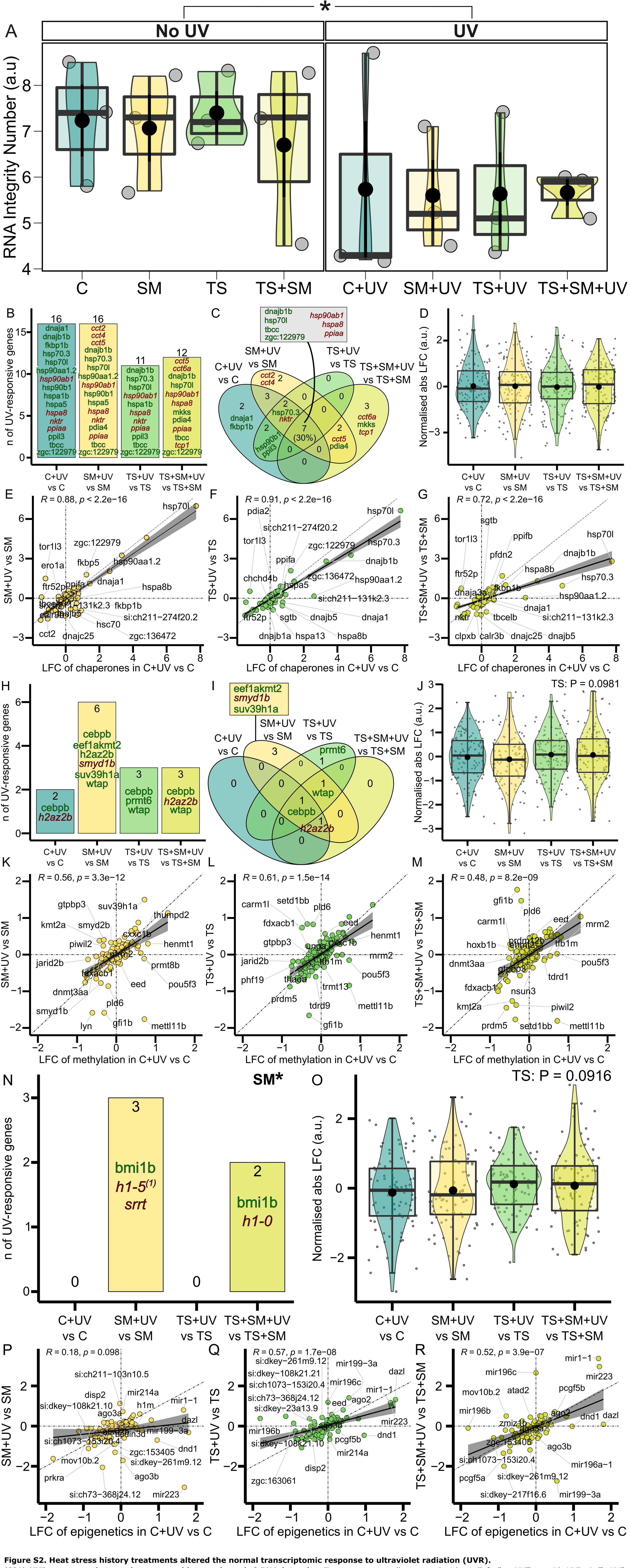


**Figure S2. Heat stress history treatments altered the normal transcriptomic response to ultraviolet radiation (UVR).**

(S2A) **UVR exposure, but not heat stress history, degraded RNA integrity.** Treatments were all compared without (left, “no UV”) or with UVB+A (“+UV”, right). *: $p < 0.05$.

(S2B-G) **TS+SM dampened the normal expression of key protein folding genes in response to UVR** (S2B) Counts and (S2C) Venn diagram of significant genes from protein folding (GO:0006457, n = 125 genes) in each treatment in response to UVR. (S2D) Magnitude of UVR-induced protein folding gene expression change represented by the normalised absolute log-fold change (LFC). Heat x medium model terms are not significant in (S2B) (Chi-squared test) and (S2D) (Scheirer-Ray-Hare test). LFC of gene expression in response to UVR in protein folding pathway in stress history treatments: (S2E) SM+UV vs SM, (S2F) TS+UV vs TS, and (S2G) TS+SM+UV vs TS+SM (y-axes) versus control C+UV vs C (x-axes).

(S2H-M) **Social metabolites altered the methylation response to UVR.** (S2H) Counts and (S2I) Venn diagram of significant genes from methylation (GO: 0043414, n = 131 genes) in each treatment in response to UVR. (S2J) Magnitude of UV-induced methylation gene expression change represented by the normalised absolute LFC. Heat x medium model terms are not significant in (S2H) (Chi-squared test) and (S2J) (ANOVA, albeit $p < 0.1$ for heat). LFC of gene expression in response to UVR in methylation pathway in stress history treatments: (S2K) SM+UV vs SM, (S2L) TS+UV vs TS, and (S2M) TS+SM+UV vs TS+SM (y-axes) versus control C+UV vs C (x-axes).

(S2N-R) **Social metabolites activated epigenetic processes in response to UVR**. (S2N) Counts of significant genes from epigenetic processes (GO:0040029, n = 84 genes) in each treatment in response to UVR. (1) Zebrafish gene is si:ch73-368j24.12 but its human ortholog is shown. (S2O) Magnitude of UV-induced epigenetic gene expression change represented by the normalised absolute LFC. Social metabolites induced more epigenetic genes in (S2N) and heat tended to increase LFC in (S2O). LFC of gene expression in response to UVR in epigenetic pathway in stress history treatments: (S2P) SM+UV vs SM, (S2Q) TS+UV vs TS, and (S2R) TS+SM+UV vs TS+SM (y-axes) versus control C+UV vs C (x-axes).

In (S2A, S2D, S2J, and S2O), black dots show mean ± s.e.m values in arbitrary units (a.u.). Boxes show median and 25%-75% quartiles. Whiskers are min/max values within 1.5 interquartile range. Individual data are given as jittered grey circles. In (S2B-C, S2H-I, and S2N), non-italicised dark green and italicised dark red gene names respectively indicate up- and downregulation (GO-term wide p-adj < 0.05 and |FC| > 1.5). In (S2E-G, S2K-M, and S2P-R), the null hypothesis is that the UVR-induced response is similar for all treatments and follows a 1-to-1 ratio (diagonal lines). Solid black lines and shaded areas depict linear fits and 95% confidence intervals. Labels indicate the top 20 genes with the most pronounced deviation relative to C+UV vs C. Treatments were C: control in fresh medium at 27°C, SM: social metabolites at 27°C, TS: fresh medium in thermal stress, TS+SM: social metabolites in thermal stress all compared to their non-UV pairs.

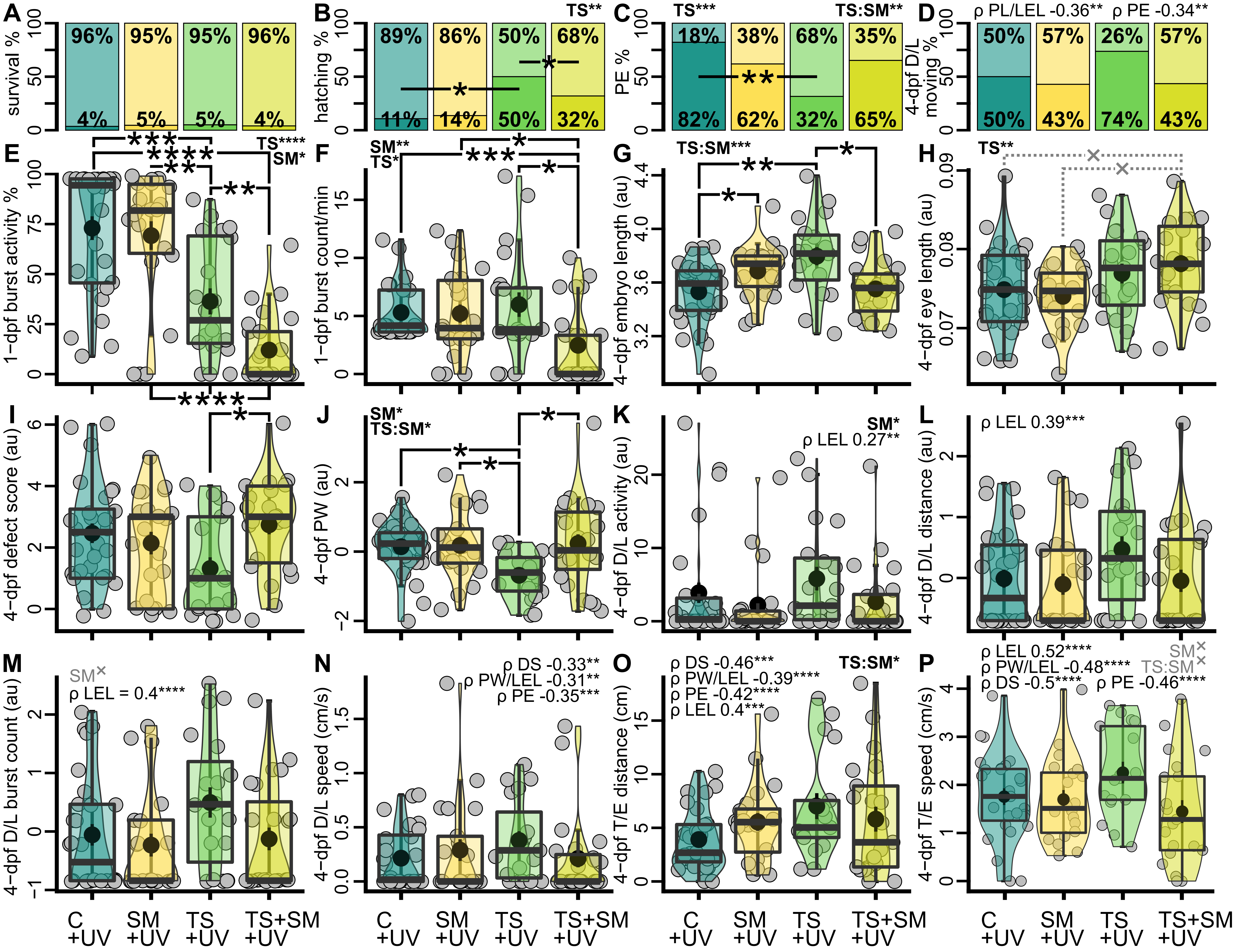


**Figure S3. Social metabolites impaired behaviour whilst heat tended to facilitate swimming through better repair.** Behaviour and fitness outcome at 1 day and 4 days post fertilisation (dpf). Values in arbitrary units (au) are transformed to fit parametric assumptions. (A) % of dead (bottom dark areas) or surviving (top light areas) 4-dpf larvae. (B) % hatched (bottom dark areas) or unhatched (top light areas) 4-dpf larvae. (C) % 4-dpf larvae with (bottom dark areas) or without (top light areas) pericardial edemas (PE). (D) % moving embryos in 4-dpf dark-light behaviour (D/L) assay. (E) Burst activity % and (F) burst count per min in response to light stimulus in 1-dpf embryos. (G) 4-dpf embryo length. (H) 4-dpf embryo size-standardised eye length. (I) 4-dpf defect score. (J) 4-dpf embryo size-standardised pericardial width (PW). (K) Swimming activity (moving time to tested time ratio), (L) total distance, (M) number of bursts accelerations, and (N) mean speed (cm/s) in the 4-dpf D/L assay. O) Total distance and (P) mean speed in the 4-dpf touch-evoked (T/E) swimming assay. Significant two-way model terms (heat x medium symbolised as "TS" and "SM") are shown in bold in top corners following analyses of deviance (binary generalised linear models) or variance (ANOVAs); or Scheirer-Ray-Hare tests. Significant correlations with covariates are shown with "ρ" symbolising Pearson's coefficient. DS: defect score, LEL: longest embryo length, PW: pericardial width, PE: pericardial edema. Pairwise comparisons are shown with horizontal bars. Treatments for phenotypic analyses were all followed by UV radiation (+UV), with C+UV: control in fresh medium at 27°C (n = 28), SM+UV: social metabolites at 27°C (n = 22), TS+UV: fresh medium in thermal stress (n = 20), TS+SM+UV: social metabolites in thermal stress (n = 25). Sample sizes are given before filtering for data with all complete cases. X: trends with weak evidence (p < 0.08, in grey), *: p ≤ 0.05, **: p ≤ 0.01, ***: p ≤ 0.001, ****: p ≤ 0.0001.

**Supplementary Dataset**

**Dataset S1** contains the results of gene expression analyses and the statistical analyses produced within this paper.
